# Thalamocortical mechanisms regulating the relationship between transient beta events and human tactile perception

**DOI:** 10.1101/2021.04.16.440210

**Authors:** Robert G. Law, Sarah Pugliese, Hyeyoung Shin, Danielle D. Sliva, Shane Lee, Samuel Neymotin, Christopher Moore, Stephanie R. Jones

## Abstract

Transient neocortical events with high spectral power in the 15–29Hz beta band are among the most reliable predictors of sensory perception. Prestimulus beta event rates in primary somatosensory cortex correlate with sensory suppression, most effectively 100–300ms before stimulus onset. However, the neural mechanisms underlying this perceptual association are unknown. We combined human magnetoencephalography (MEG) measurements with biophysical neural modeling to test potential cellular and circuit mechanisms that underlie observed correlations between prestimulus beta events and tactile detection. Extending prior studies, we found that simulated bursts from higher-order, non-lemniscal thalamus were sufficient to drive beta event generation and to recruit slow supragranular inhibition acting on a 300ms time scale to suppress sensory information. Further analysis showed that the same beta generating mechanism can lead to facilitated perception for a brief period when beta events occur simultaneously with tactile stimulation before inhibition is recruited. These findings were supported by close agreement between model-derived predictions and empirical MEG data. The post-event suppressive mechanism explains an array of studies that associate beta with decreased processing, while the during-event faciliatory mechanism may demand a reinterpretation of the role of beta events in the context of coincident timing.

---

While considerations of regular, sustained oscillations have supported many theories of neural encoding (e.g. Kay et al. 2009; Fries 2015), neural oscillations often appear only briefly, raising subtle questions as to precisely if, and how, a transient “rhythm” can pertain to sensory and cognitive processing (Jones 2016; Cole and Voytek 2017; van Ede et al. 2018). The observation that brain dynamics are nonstationary (e.g. Jestrović et al. 2014) — with time-localized spectral energy on individual trials non-uniformly contributing to time-averaged spectral power — has become a focus of a number of research efforts (Xing et al. 2009; Jones et al., 2009; Feingold et al. 2015; Jones 2016; Lundqvist et al. 2016; Cole and Voytek 2017; Shin et al. 2017; van Ede et al. 2018, De Gennaro and Ferrara 2003)). Modern consensus holds that many brain rhythms tend to occur as sparse, energetically concentrated *events* (cf. *bursts*^*^) that can have marked influence on waking perception and behavior.

Transient beta band (15–29Hz) rhythms are one of the most well-established *event-like* rhythms, observed in sensory (Sherman et al. 2016; Shin et al. 2017), motor (Rule et al. 2017; Little et al. 2018, Wessel 2020), prefrontal (Lundqvist et al. 2016; Sherman et al. 2016; Lundqvist et al. 2018), and subcortical regions (Feingold et al. 2015), and across species and recording modalities (Shin et al. 2017). Modulation of beta events, as quantified in event rate or time-dependent probability density, are associated with memory processes (Lundqvist et al. 2016; Lundqvist et al. 2018), sensory perception (Shin et al. 2017) and motor action in both health and disease (Torrecillos et al. 2015; Cole et al. 2017; Tinkhauser et al. 2018; Little et al. 2018, Wessel 2020). The neural mechanisms by which beta events relate to behavior are currently unknown, and their cognitive and perceptual roles remain in debate (Engel and Fries 2010; Jenkinson and Brown 2011; Spitzer and Haegens 2017).

We have observed that the rate and timing of beta events in magnetoencephalographic (MEG) and local field potential (LFP) recordings from primary somatosensory cortex (SI) predict threshold-level tactile perception (Sherman et al. 2016; Shin et al. 2017). Two or more events in a one-second prestimulus period decreased perception, with perception impaired for 100–300ms after a beta event (Shin et al. 2017). Here, our goal was to uncover the neural mechanisms underlying this relationship between beta events and tactile perception. To this end, we applied computational neural modeling designed to interpret the circuit mechanisms of human MEG (or EEG) signals based on their biophysical origin (Neymotin et al 2020). In prior research, we applied this modeling framework to study the neural mechanisms underlying prestimulus SI beta events (Jones et al 2009; Ziegler et al 2010; Sherman et al 2016) and of post-stimulus tactile evoked responses (Jones et al 2007; Jones et al 2009). Here, we join and extend these prior studies.

We first examine the relationship between prestimulus beta events and evoked correlates of tactile detection in our MEG data and then combine simulation of beta events and tactile evoked responses to interpret neural mechanisms mediating this relationship. Our prior study showed that beta events are generated by synchronous bursts of excitatory synaptic drive to infragranular and supragranular cortical layers, where the supragranular drive is stronger and lasts one beta period (~50ms) (Sherman et al 2016). The supragranular drive excites distal apical dendrites of pyramidal neurons inducing downward dendritic currents to produce a trough of ~50ms duration, which is the dominant feature of recorded beta events. The bursts of activity generating this supragranular drive are thought to originate in “higher-order” (cf. “modulatory”, “nonlemniscal”, or “matrix”) thalamus (Mo and Sherman, 2019, Jones, 1998; Jones 2001; Sherman 2016). We hypothesized further that slow inhibitory currents at the GABA_B1a_ timescale (Otis et al. 1993) are also recruited by these distal-targeting bursts, based on the observation that inhibitory effects of beta events on tactile detection last ~100-300ms (Shin et al 2017). By extending our model to incorporate this hypothesis, we were able to account for the observed relationship between prestimulus beta events and evoked correlates of tactile detection, providing a mechanistic explanation as to why beta events often precede non-detected tactile stimuli. Our simulations predicted -- and further data analysis revealed -- that the exogenous thalamic drive that generates beta events also creates a brief window of excitation: Tactile evoked responses occurring during beta events are enhanced, and perception is facilitated, before the long-timescale inhibition is engaged. The model suggests the beta associated sensory facilitation and suppression come down to the biophysical ability of SI pyramidal neurons to spike in response to tactile stimulation, and hence to relay sensory information to other parts of the brain. These results are discussed in relation to an array of studies that associate beta with decreased processing, as well as literature that suggests beta may reflect timed prediction and learning of cognitively relevant stimuli.

## Materials and Methods

### MEG data collection and analysis

We analyze data from a previous study, where MEG data localized to primary somatosensory (SI) cortex were collected during a tactile detection task (Jones et al. 2007). Detailed MEG and behavioral data collection methods can be found in the original study (Jones et al. 2007), and in a follow up study that analyzed the same data (Jones et al. 2009). Relevant details and data analysis specific to the current study are described below.

#### Tactile-detection task

Subject’s (N=10 18–45-year-old adults; 6 female) performed a tactile detection task during whole-head 306 channel MEG recordings (Elekta Neuromag Vectorview). Data were sampled at 600Hz and bandpassed online between 0.01Hz and 200Hz. Subjects were seated with eyes open and focused on a fixation cross in the middle of screen. Each 3 second trial began with a 2kHz auditory cue presented to both ears for a duration of 2000ms. A tactile brief tactile stimulus was delivered at a time sampled uniformly from a comb distribution spanning 500–1500ms after cue onset, consisting of 11 evenly spaced postcue timings. The stimulus consisted of a single sinusoidal tactile pulse lasting 10ms, delivered from a piezoelectric ceramic benderplate to the tip of the D3 digit of the right hand. After auditory cue offset, subjects were instructed to press a button with the left hand (second or third digit) to report whether the stimulus was perceived. Tactile stimulus pulse amplitude was initially tuned to the individual’s perceptual threshold using the parameter estimation by sequential testing method (Dai 1995; Leek 2001) and then dynamically maintained to a 50% detection threshold (Jones et al. 2007). For each individual, the final 100 hit (detected) and 100 miss (non-detected) trials were isolated for further analysis due to the stability of the dynamics at the end of the experiment (Wan et al. 2011). In the current study, data were analyzed from 1 second before to 140ms after the tactile stimulus.

#### MEG current dipole source localization

Activity from the hand-region of SI was localized using a two-dipole model (SI and SII), fitting suprathreshold responses with a signal-space projection method (Tesche et al. 1995; Uusitalo and Ilmoniemi 1997), see Jones et al 2007 for details. The localized SI signal analyzed in the current study represents a **primary current dipole**, with units of current x distance (Ampere-meters, Am). The neural model used in this study, detailed below, was specifically designed to simulate the primary current dipole signals, with directly comparable Am units, as in the Human Neocortical Neurosolver software (Jones et al. 2009, Neymotin et al 2020).

#### Defining beta events and assessing their perceptual effects

Events in the prestimulus period were identified as regions in the time-frequency plane whose energy crossed six times the median energy at a given frequency, computed separately for each individual by convolution with a six-cycle Morlet wavelet (see Shin, et al 2017 for more detailed methods). In this study we restricted our analysis to events crossing this threshold between 20–22Hz, following the spectral concentration within the 15–29Hz beta wideband observed in human somatosensory recordings (Jones et al. 2009; Sacchet et al. 2015). Trials were then partitioned into those with one or more prestimulus events (*event* trials) and those without events (*no-event* trials), after which we reverified that these narrowband beta events affected perception via the Wilcoxon test.

#### Evoked response time-derivative estimation

During evoked responses, bias may be introduced by beta events in the prestimulus period (see e.g. Iemi et al, 2019), and so we did not baseline normalize the evoked responses in this study. Instead, we examined both raw data and time-derivatives — these yield baseline-free comparisons of dipole changes from active membrane processes. We used total-variation regularized differentiation, which corrects noise amplification in first-difference estimates of time-derivatives while preserving jump discontinuities (Chartrand 2011) observed in MEG data, without linear filtering artifacts. We fixed the regularization parameter at 10^−8^ based on visual inspection of regularized fits to two trials.

#### Post-stimulus phase coherence

For post-stimulus intertrial phase-coherence analyses, we first filtered the data in the frequency band of interest. To assure robustness of the results, we used both Chebyshev and Butterworth filters in nearby frequency bands. For the phase coherence analysis over the low frequency 18-24Hz band in Figure 10B, results from a 3^rd^ order Butterworth filter are reported (a 20-22Hz Chebyshev filter with 10dB within-band ripple gave similar results), while for the phase coherence over the higher frequency 160-200Hz band in Figure 10G/H, we used a Chebyshev filter with 10dB within-band ripple (a 3^rd^ order Butterworth filter at 178--197Hz yielded similar results, but higher-order Butterworth filters were numerically unstable). The same Chebyshev filter was applied to both model and MEG data in Figure 10H. After bandpassing, we computed the (*n.b.* nonlocal) Hilbert transform for each trial, computed the complex unit vectors associated with each timepoint (with phase given by the complex angle of the unit vector), and then found cross-trial phase coherence as the modulus of the complex mean of unit vectors over trials.

### Computational Neural Model Construction and Analysis

#### SI circuit model

We adapted the model structure from Jones et al. 2009 to reflect additional anatomical and physiological detail. Several changes to the 2009 model’s free parameters led to improved data fits and were informed by use of the Human Neocortical Neurosolver software (Neymotin et al. 2020). HNN freely distributes all of the code from the 2009 model, from which the current parameters were adapted. A description of differences between Jones et al. 2009 and the current model is in Supplementary Materials and Methods, and a direct comparison between parameters in each study are in Supplementary Table 1.

In brief, the model represents a layered cortical column with multicompartment pyramidal neurons and single compartment inhibitory neurons in supragranular layers 2/3 (L2/3) and infragranular layer 5 (L5), synaptically coupled with glutamatergic and GABAergic synapses (Figure 1).

**Figure 1.**
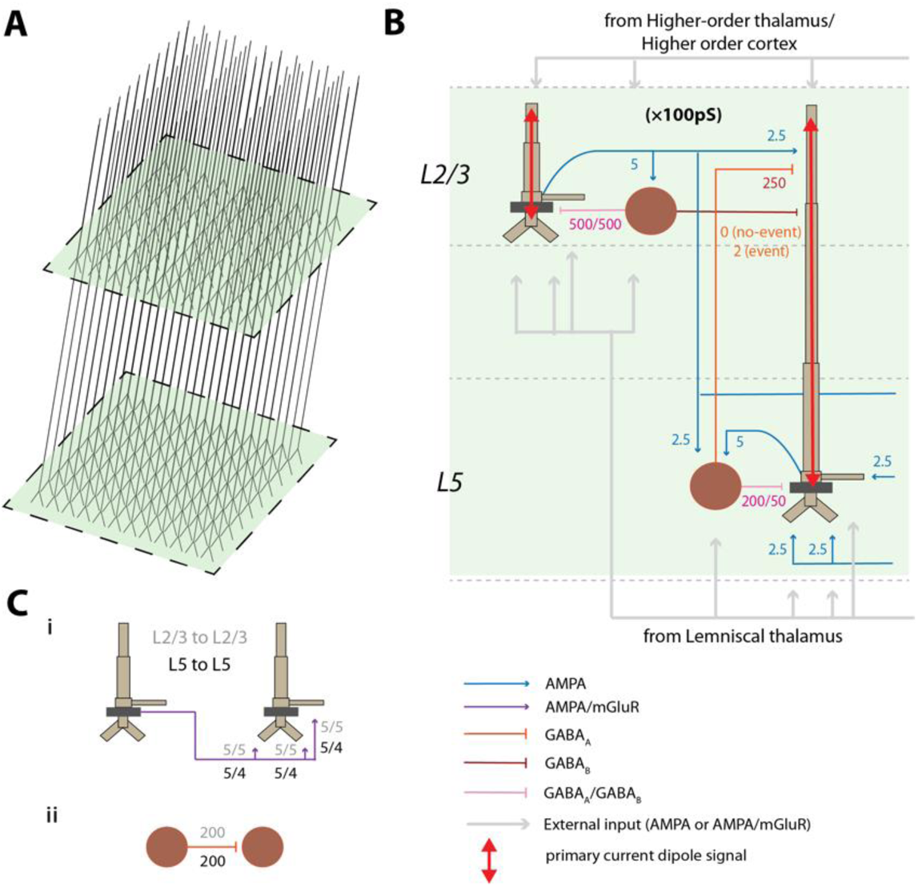
SI computational neural model. **A)** Model 3D pyramidal geometry shown approximately to scale. **B)** Reduced circuit schematic, with somata indicated in darker brown. Maximal conductances for each synaptic site are given in multiples of 10^−5^ nS/cm^2^. Red arrows indicate the source of the primary current dipole calculated from the intracellular current flow in the pyramidal neuron dendrites. The units of measure of this signal are directly comparable to MEG data (Am). **C) i)** Schematic of lateral connections between pyramidal cells with synaptic weights labeled. **ii)** Schematic of lateral connections among interneurons.

The MEG signal studied is generated by a **primary current dipole** localized to SI. As described in detail in prior studies (Jones et al. 2007; Jones et al. 2009; Neymotin et al. 2020), the primary current dipole signal is simulated by net intracellular current flow in the pyramidal neural dendrites in the direction parallel to the apical dendrites, multiplied by the length of the cell, and summed across all pyramidal neurons (see red arrows in Figure 1B). These intracellular currents are generated by the interaction of active and passive membrane properties and synaptic dynamics. Synaptic strengths in the current model are detailed in Figures and all other parameters used are detailed in Supplementary Table 1. The units of measure of the simulated primary current signal are directly comparable to the source localized MEG data (current x distance, Am), where a scaling factor is applied to the net response to match the amplitude of the recorded signals and provides an estimate of the size of the network contributing to the recorded response, described further below.

Details of how the evoked responses and beta events were simulated in the model are described in the Results section. Further details are in Supplemental Computational Model Methods, and a discussion of modeling limitations and parameter robustness are in the Discussion section.

### Comparing MEG and Model Evoked Responses

Our initial analysis of the SI MEG tactile evoked responses relied on finding time-localized differences between event and no-event trials, and between hit and miss trials (see Figure 3), which we used to focus the model examination. The statistical methods for this comparison, and for the validation of the model-derived prediction on phase coherence among evoked responses (Figure 10), are described here. In both cases, because the number of statistical tests is equal to the number of timepoints, multiple comparisons correction is accounted for by adopting a modified max-t test to limit familywise error rate, as follows:

**Figure 3:**
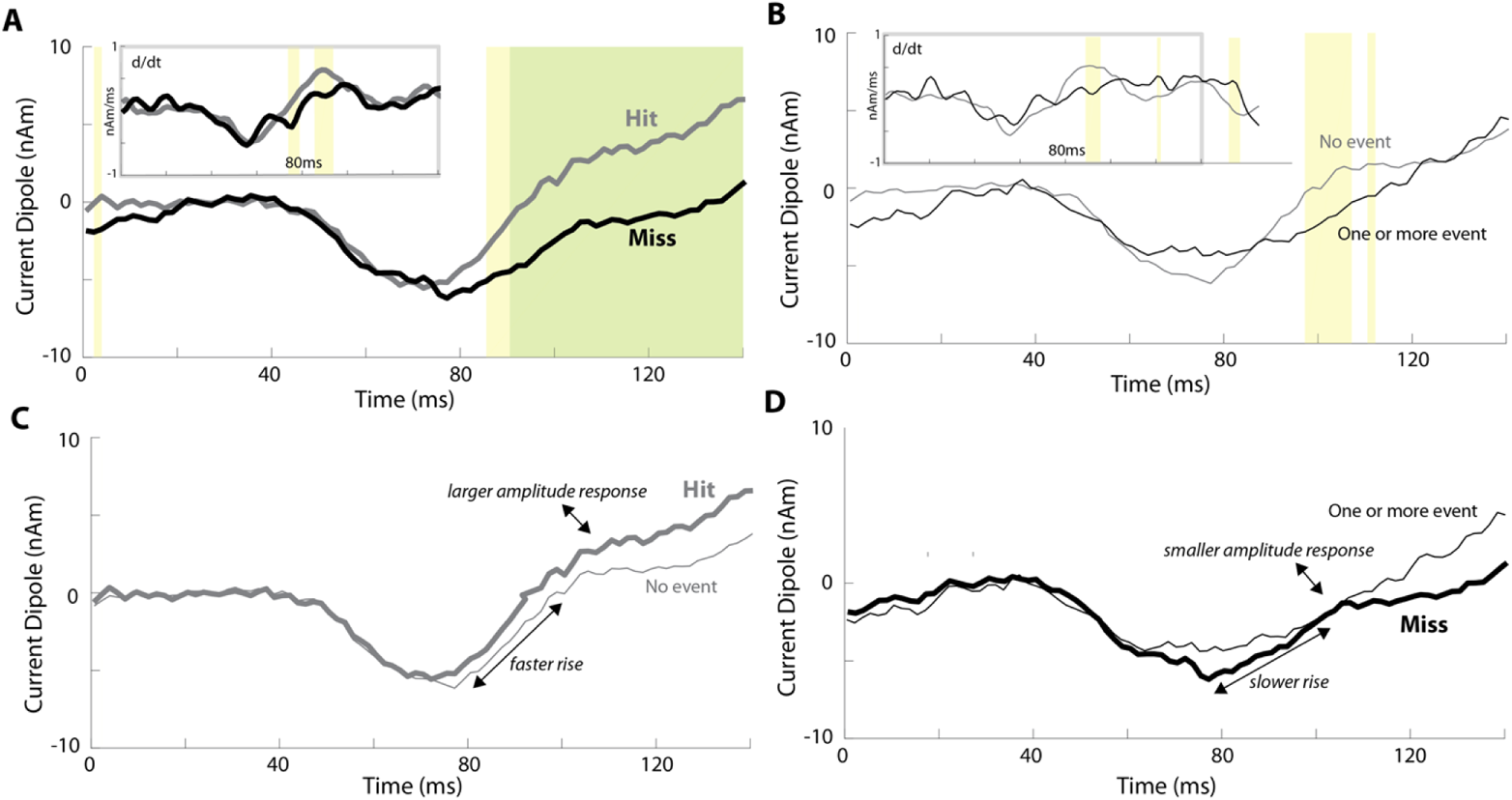
Human tactile evoked responses conditioned on prestimulus beta events and perception. **A)** Mean evoked responses for detected (hit) and nondetected (miss) trials. **B)** Mean evoked responses for trials with (one or more events) or without (no event) a prestimulus beta event. Significance is reported both pointwise in yellow (p < 0.05; permutation test) and after FWER-correction in green (α = 0.05; modified max-t test; see Methods). *Insets:* Mean evoked response time-derivatives (after total-variation regularization; see Methods). **C)** Overlay of hit and no event trials. **D)** Overlay of miss and one or more event trials.

**Figure 10:**
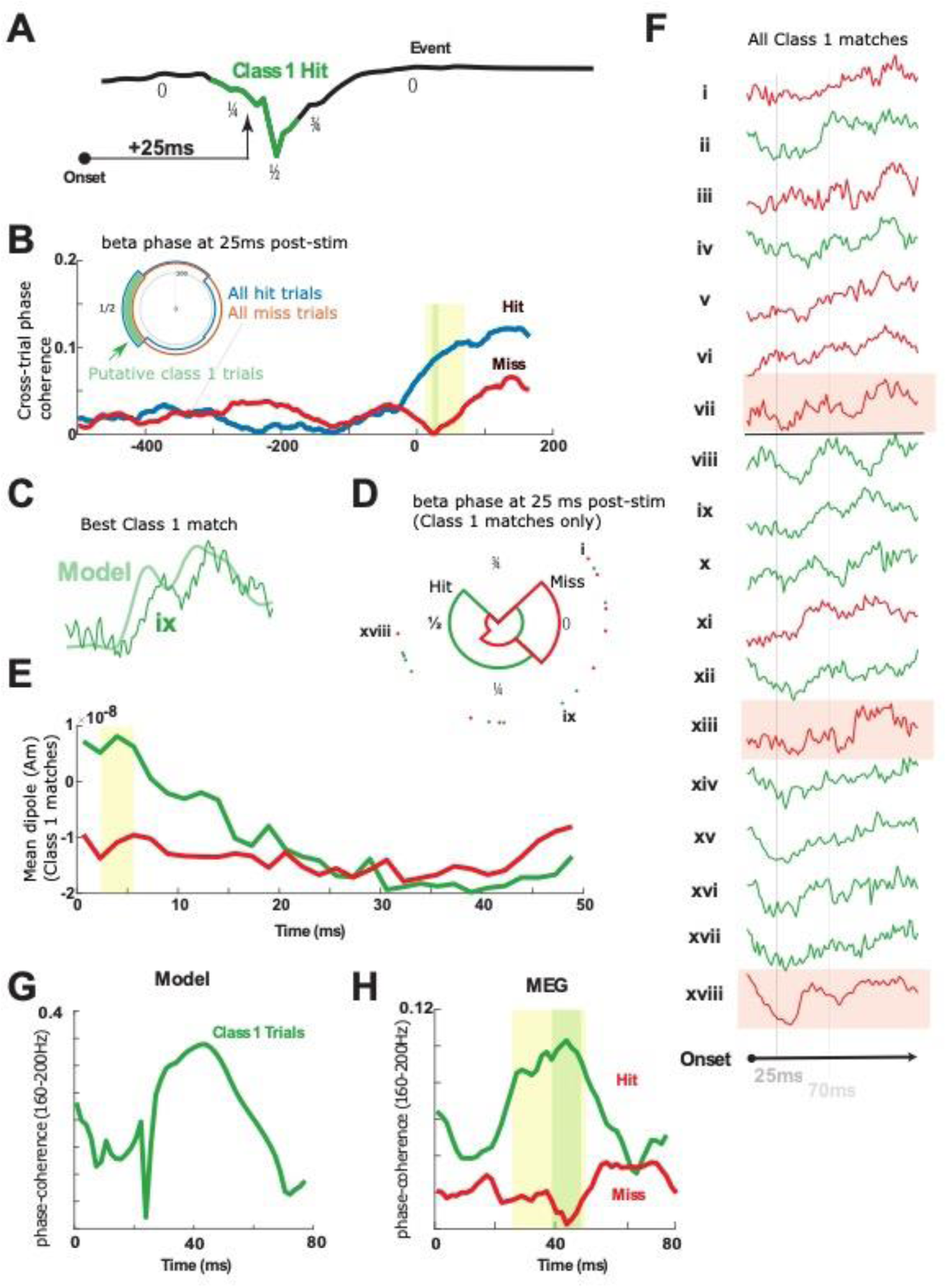
Model Class 1 responses correlate with hit trials. **A)** Model beta event phases. Green region depicts the approximate phase interval where Class 1 trials are predicted (phase interval 0.25 – 0.75; normalized) given the 25ms conduction delay from the periphery. **B)** Across-trial beta phase coherence (18-24Hz) in the MEG evoked response is shown for hit (blue) and miss (red) trials. Poststimulus beta coherence is higher in hit trials at precisely 25ms. *Inset*: Phase histogram of beta phase at 25ms post-stimulus on hit and miss trials. Putative Class 1 hit trials are shown in green. **C)** Example correlation between the model Class 1 response and empirical MEG evoked response during a hit trial. **D)** Beta phase at 25ms poststimulus nearly completely separates the Class 1 correlates into hit and miss trials (18 highest correlation trials; 4 bins). Hit trials occurred within the phase interval predicted by the model with one exception. **E)** The eighteen trials with highest correlation to Class 1 response averaged separately over hit and miss trials, show that hit trials exhibit an initial downward dipole slope. **F)** Direct inspection of all high correlation Class 1 responses shows that 10 of 10 hit trials were either in dipole troughs or in downward phases at 25ms poststimulus, while 5 of 8 miss trials are in rising phases or plateaus; 3 outliers shaded pink. **G**) Across-trial high frequency coherence (160-200Hz) in model Class 1 responses predicts a peak in coherence near 40ms post-stimulus in hit trials. **H)** Across-trial 160–200Hz coherence distinguishes hit from miss trials near 40ms poststimulus (1000 hit trials, 1000 miss trials) in precise agreement with the model prediction. In all panels, yellow shading indicates p < 0.05 (pointwise permutation test), and green shading indicates α = 0.05 (FWER-corrected; modified max-t test; see Methods). Significance in panel B is preserved at α = 0.02, and significance in panel H is preserved at α = 0.01.

Let *X* be a data matrix with *X*(*i, t*) representing the observation at time *t* during trial *i*. Here *X*(*i, t*) will be some property, such as current dipole or instantaneous phase, of a MEG signal from a single source-localized channel (real or modeled) at time *t* on trial *i*. Let *L* be vector of labels with *L*(*i*) representing the label for trial *i*. Here *L* ∈ {0,1} will indicate a behavioral outcome or prestimulus event, i.e. a correct detection or a beta event, for trial *i*.

We use permutation tests to test the null hypothesis that the labels and the data at time *t* are independent (or, more generally, that the labels are exchangeable given the data, at time *t*). For current dipole data our test statistic is the difference between the average current dipole on trials with label 1 and the average current dipole on trials with label 0. For instantaneous phase data our test statistic is the difference between the phase coherence on trials with label 1 and the phase coherence on trials with label 0.

Permutation tests work by randomly shuffling the trial labels and recomputing the test statistic for nonsensically labelled data many times in order to obtain a null distribution. The observed test statistic (on the correct trial labels) is compared to this null distribution in order to obtain a p-value. We report two-sided p-values in all cases and use at least 5000 permutations. See (Lehmann and Romano 2006) for more details about hypothesis testing and permutation tests. We use the same trial label permutation for testing each time *t* (i.e. the shuffling is performed on entire time-series, not independent time points). This is important for our method of controlling for multiple hypothesis tests, described next.

Because we are testing a separate null hypothesis at each time *t*, we controlled for multiple hypothesis tests using a variant of the max-t method, which creates a global test statistic by taking the maximum (for the upper-tail) and minimum (for the lower tail) of the test-statistic over all times *t*, and then creates a null distribution (one for each tail) in the usual way by shuffling trial labels. The observed test statistic at each time *t* is compared to these common, global null distributions in order to create adjusted p-values that provide strong control of the family-wise error rate. Rejecting those null hypotheses with adjusted p-values ≤ α guarantees that the probability of zero false rejections is ≥ 1 − α. See (Westfall and Young 1993; Nichols and Hayasaka 2003) for more details about permutation tests and multiple testing.

Our variant of the max-t method robustly standardizes the test statistics at each time *t* prior to computing the maximum (or minimum) in order to more evenly distribute statistical power across all of the hypotheses. The specific details of our procedure can be found in (Amarasingham et al. 2011 Nov 18).

## Results

### Human MEG reveals a relationship between prestimulus SI beta events and tactile detection

#### A Single Narrowband Prestimulus Beta Event is a Signature of Decreased Tactile Detection

We had previously shown in human MEG and mouse LFP that two or more prestimulus 15–29Hz SI beta events yield a bias toward non-detection of a perceptual threshold-level tactile stimulus (Shin et al. 2017). Beta events were defined as periods with power above a 6x-median threshold in the time-frequency plane and typically lasted <150ms (see Materials and Methods; Figure 2A). Though the presence of a single 15–29Hz event did not significantly influence detection (Shin et al. 2017), we observed that human beta events tend to concentrate in a narrowband near 20Hz (e.g. Figure 2A; note that in the figure, power is above 6x-median in the 20–22Hz band but peaks near 18Hz). When restricting our analysis to the 20-22Hz range, trials with one or more prestimulus beta event (*event* trials) were associated with lower detection probabilities compared to *no-event* trials (8/10 participants Figure 2B, p < 0.01; Wilcoxon rank-sum test; in total: 502 event trials (295 miss, 207 hit) and 1498 no-event trials (705 miss, 793 hit); per-subject mean and standard deviation 50.2 ± 9.5 event trials, 148.8 ± 9.5 no-event trial; median reduction 10%; maximum reduction 28%).

**Figure 2:**
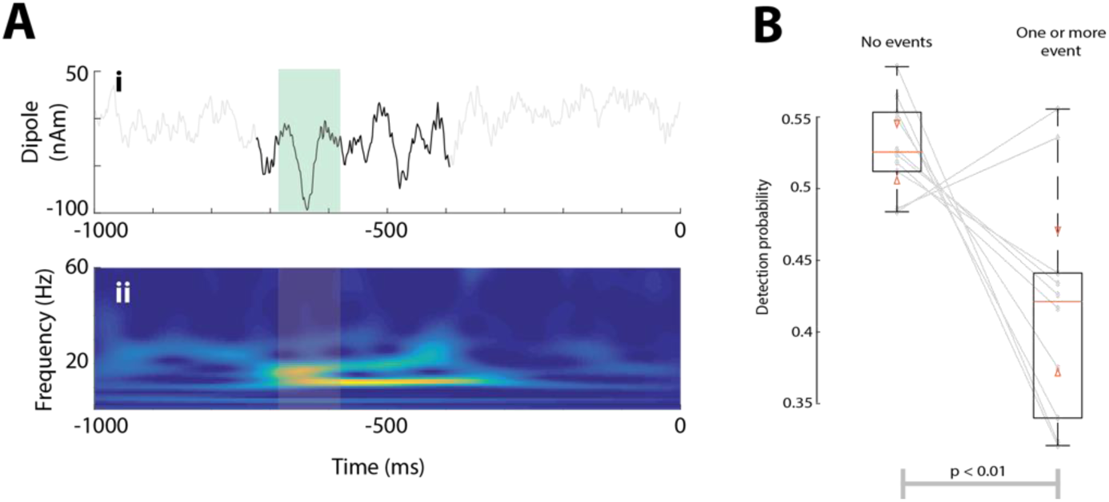
Prestimulus beta events are associated with lower near-threshold detection probabilities in human SI. **A)** Example trial with a beta event in **i)** the time-domain with beta event in black (stereotypical beta cycle is highlighted in green) and **ii)** as a wavelet spectrogram with event timespan highlighted. **B)** Detection probabilities for all participants. Boxes show interquartile ranges. Orange bars and triangles represent median and comparison intervals respectively; see Methods. A beta event with > 6 median power at 20—22Hz in the prestimulus period reduces detection probability (p < 0.01; Wilcoxon rank-sum test) by approximately 10% on average, and on an individual basis a prestimulus beta event alters detection probability in a range of −28% to +7%.

#### Amplitude and slope differences in the 80-110ms post-stimulus time period link beta events and correlates of detection in the tactile evoked response

Our prior studies showed that the tactile evoked waveform has a slower rise and smaller amplitude after the earliest prominent trough at ~70ms on non-detected trials (Jones et al. 2007). These results are refined in Figure 3A, where the regularized time derivative of the signal differed between detected (hit) and non-detected (miss) trials at ~75-97ms post-stimulus (Figure 3A; inset), with smaller derivatives, and smaller waveform amplitudes from ~87-140ms on miss trials (Figure 3A; yellow: p < 0.05; permutation test, uncorrected; green: alpha = 0.05; FWER-corrected permutation test)). We next set out to understand if prestimulus beta events influence the evoked response in a similar manner, indicating a potential causal relationship between beta generating mechanisms and detection. We found that the two perceptual effects described above were also observed when comparing prestimulus event and no-event trials (Figure 3B). Prestimulus events led to a slower rise ~80-95ms post-stimulus, (Figure 3B inset), and to smaller evoked amplitude from ~98-113ms (Figure 3B; yellow bars; p < 0.05; permutation test, uncorrected). Figures 3C and 3D highlight the similarities further by plotting the no event and hit trials together, and the one or more events and miss trials together.

We note that the due to the small amplitude threshold level stimulus used in this study, earlier components of the evoked response (< 70ms) are difficult to distinguish in the macroscale current dipole signals, and circuit difference may not be visible in the recorded MEG signal. However, our modeling results below showed clear differences at the cellular level on trials with and without beta events beginning at ~25ms post-stimulus. We further note that the lower amplitude baseline at time zero on miss/beta event trials (Figure 3A/B) is consistent with the assumed beta generation mechanisms which creates downward deflecting SI current dipoles (see Figure 5 below).

**Figure 5:**
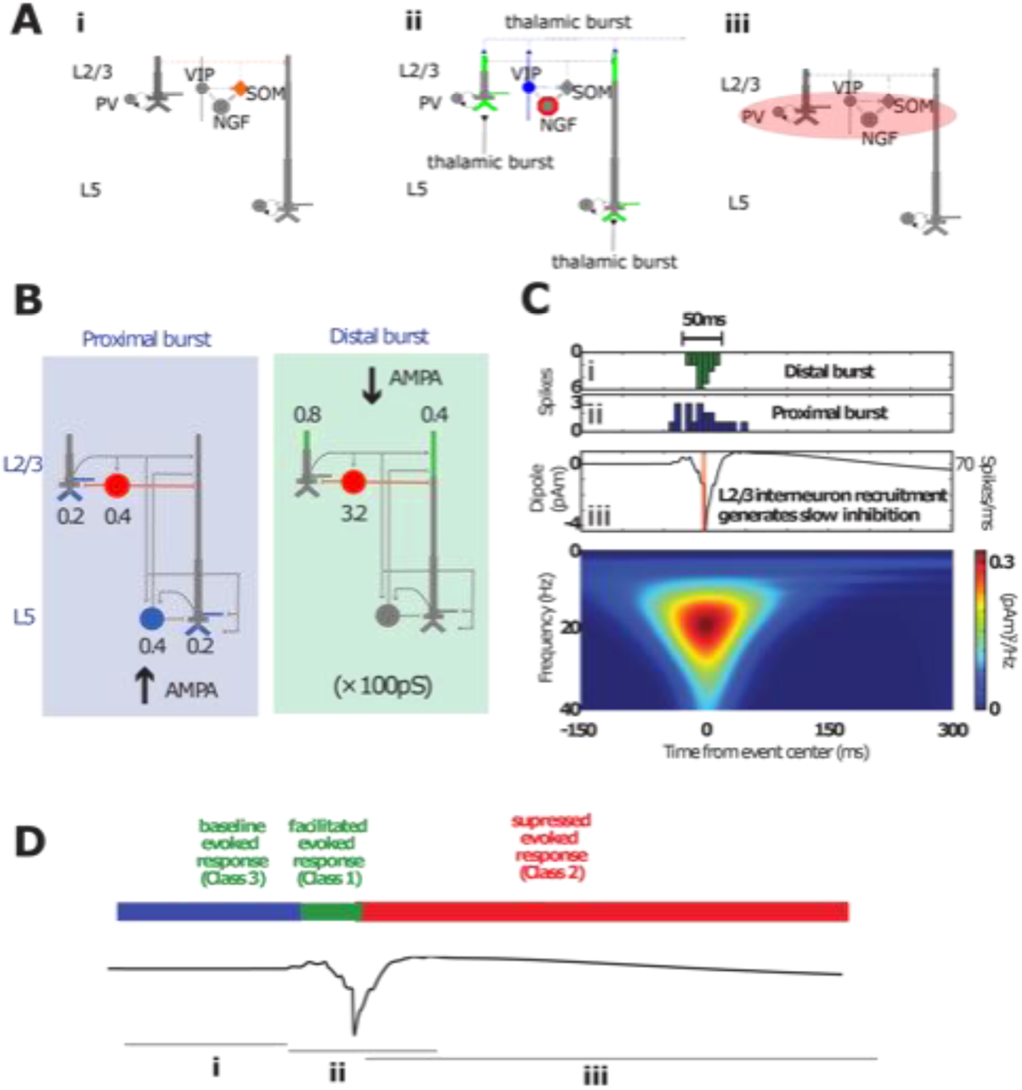
Simulating beta events and the associated recruitment of inhibition in the SI model. **A)** Idealization of the SI circuit activation pattern before, during and after a beta event. **i)** Before an event, SOM+ cells are active, inhibiting the pyramidal dendritic tuft. **ii)** During an event, a thalamic burst induces the VIP+ activation that inhibits SOM+ and activates NGFs via gap junctions, and also excites tufts of pyramidal dendrites, generating a downward current dipole. **iii)** Neurogliaform cells activate metabotropic GABA_B_ receptors, with a delay in inhibition due to their intrinsic dynamics and the slow rise time of GABA_B_ currents. Individual interneuron types are not explicitly modeled. The essential model features is simulation of interneuron firing that generates GABA_B_ currents at the time of the beta event trough. **B)** Model of incident thalamocortical burst through proximal and distal project pathways, with weights onto cell compartments specified. **C)** Model implementation of the thalamic burst generating a beta event **i).** Histogram of spikes incident on the distal (green) and proximal (**ii**, blue) dendrites of the cortical pyramidal and inhibitory neurons, as shown in (B), which generate the simulated beta event. **iii)** Corresponding current dipoles with concurrent spike histogram. Only the model L2/3 inhibitory neurons fire action potentials during a beta event. Activation of these interneurons causes GABA_B1a_ inhibition on pyramidal neurons, as shown in (B; red). **iv)** Time-frequency representation of the simulated beta event. The resulting time-dependent modulation in circuit sensitivity to tactile evoked input is schematized in **(D)**.

### Computational modeling shows burst mechanisms producing SI beta events also generate the observed relationship between beta events and correlates of detection in the tactile evoked response

Having established an empirical relationship between prestimulus beta events and tactile detection (Figure 3), we next sought to understand circuit mechanisms by which this could occur using our computational neural modeling framework. We begin with a review of how evoked responses and beta events are reproduced in the model and then simulate them together.

#### Simulating a tactile evoked response

To model the tactile-evoked response, we simulated a sequence of external drives to the local cortical network through layer-specific pathways based on known sensory evoked inputs to SI, as in our prior studies (Jones et al. 2007; Jones et al. 2009; Ziegler et al 2010; Sliva et al 2017; Neymotin et al 2020) and reviewed here (Figure 4A, Supplementary Table 1). Sensory input first arrives from the periphery through the lemniscal thalamus to granular layers at 25ms and then propagates directly to the supragranular and infragranular layers. This initial cortical input is simulated with a “proximal” or “feedforward” synaptic drive to the local network (Figure 4A, left).

**Figure 4:**
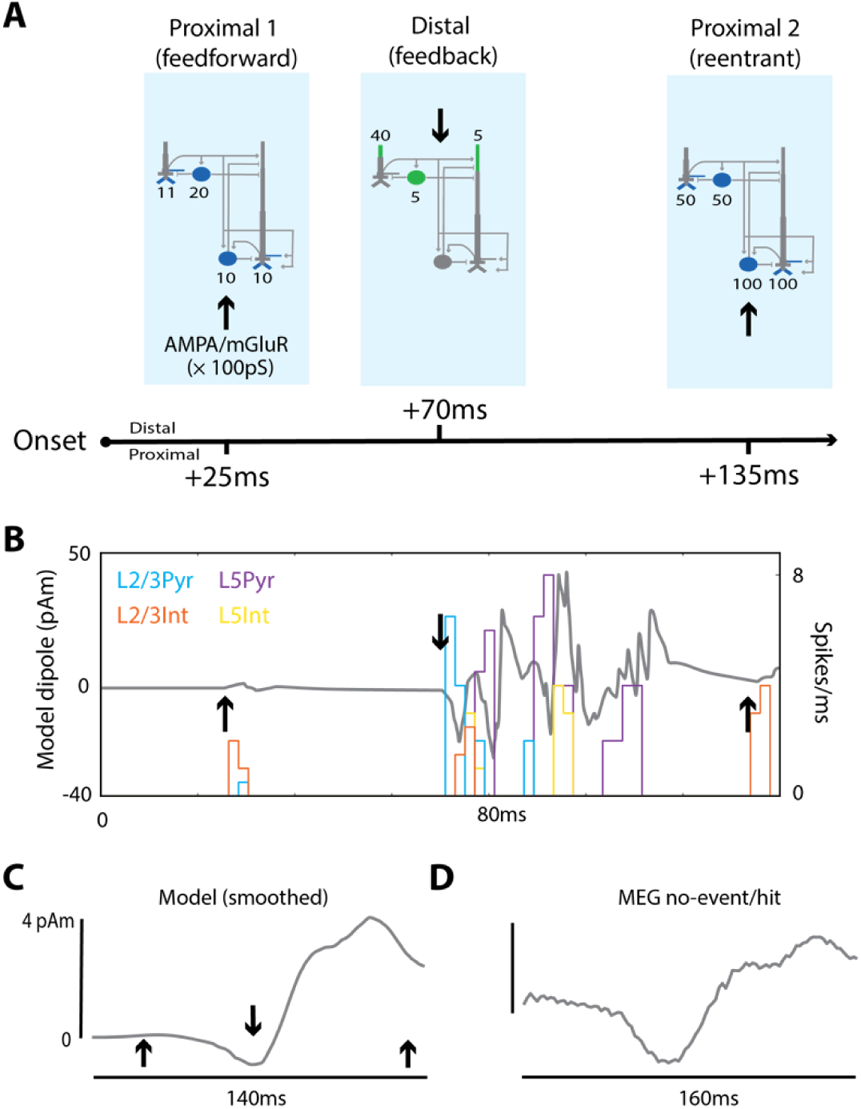
The SI model qualitatively reproduces the MEG-measured SI no-event/hit evoked response. **A)** Schematic of the feedforward/feedback exogenous input sequence reproducing the somatosensory evoked response. Sites of proximal glutamatergic “feedforward” input are shown in blue and sites of distal “feedback” inputs in green. Pyramidal cells have three proximal sites (two on the basal dendrites, one on the oblique dendrite), each with the weight indicated here. **B)** Spike histograms and raw sensory evoked response from the model at rest (i.e. without a prestimulus beta event): arrows same as in (A). **C)** The same model evoked response after smoothing by convolving with a 45ms Hamming window. **D)** MEG evoked response averaged over all hit trials without a prestimulus beta event.

To reflect the threshold nature of the task, the strength of this feedforward drive was tuned to create minimal spiking in the L2/3 cells, and to not cause L5 cells to spike. Spiking in the L2/3 pyramidal neurons then creates backpropagating action potentials and generates a small positive deflection in the current dipole response (recall that the size of the response is proportional to the length of the dendrites). This is followed by recurrent inhibition at the soma that pulls current down the dendrites and generates a small negative deflection. Note that, while this early response is observed in the model, it is obscured in the macroscale MEG data (Figure 3) which represents activity averaged over a larger network, as discussed further below.

At 70ms, excitatory feedback synaptic inputs — likely originating in SII (Cauller and Kulics 1991) — arrive at supragranular targets, activating L2/3 interneurons and L2/3 and L5 pyramidal neuron tufts (see distal /feedback synaptic drive in Figure 4A; middle). This drive initially pushes current flow down the pyramidal neuron dendrites in both layers to generate a negative deflection near 70ms. Subsequently, the pyramidal neurons in both layers generate persistent action potentials (via local recurrent synaptic excitation) that propagates up their apical dendrites to induce a positive deflection between ~80-120ms. At 135ms, a second, stronger “feedforward” input is presumed to arrive as part of an induced thalamocortical loop of activity (see proximal 2 /re-entrant synaptic drive in Figure 4A right). This drive once again generates spiking in the pyramidal neurons, that propagates up the dendrites to induce a late positive deflection near 140ms. The raw SI current dipole response (gray) and net spiking activity in each cell population during this sequence of external perturbations are shown in Figure 4B, where black arrows mark the times of the exogenous drive. A smoothed version and comparison to the MEG measured SI evoked response are shown in Figure 4C/D. This smoothing account for spatiotemporal averaging that occurs when recording the response over a large heterogeneous network in the data as compared to the model.

Importantly, in tuning the model parameters, we began with the evoked response parameters used in our prior modeling study (Jones et al 2009). We then adjusted the evoked response proximal and distal input parameters until a single set of input parameters could produce both the event and no event waveforms as observed in our data, particularly the difference between event and no event cases near ~100ms, as in Figure 3B (see comparison of parameters used to those in Jones et al 2009; Supplementary Table 1). This single set of input parameters was applied in all simulations; *post hoc* model tuning was not performed for quantitative data fits (Figure 9) or predictions in subsequent analyses (Figure 10).

**Figure 9:**
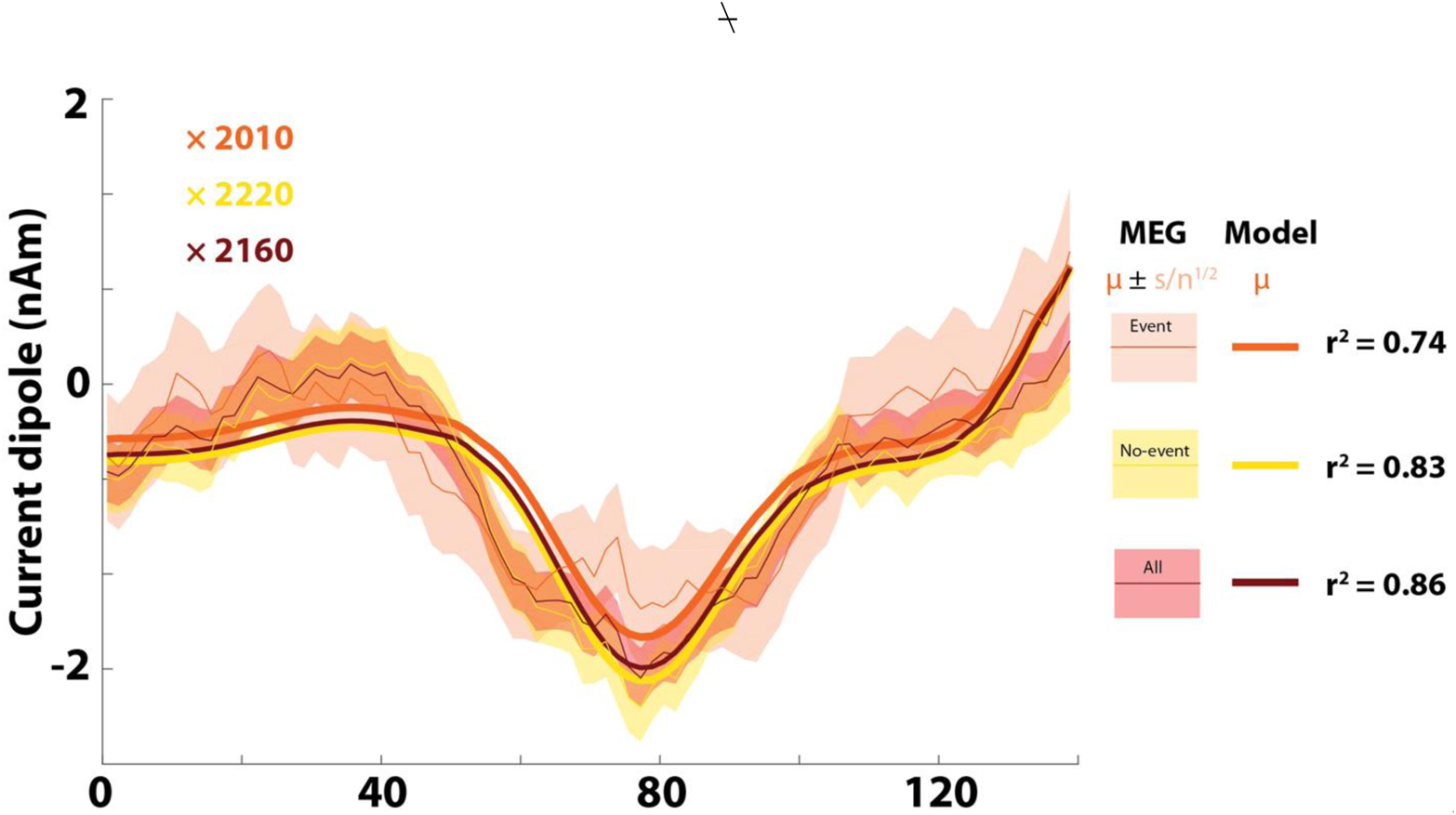
Model Class 2 responses correlate with miss trials. **A.** The model Class 2 response directly correlates with the average MEG response for miss trials, with or without a prestimulus beta event. The legend (right) indicates the model (thick lines) and MEG data distributions (thin lines; shading indicates standard error of the mean) for trials with and without events, and for all trials in aggregate. The model was fit by regressing directly onto the empirical means, and the corresponding r^2^ values and fitting parameters confirm a consistent agreement between model Class 2 trials and miss trials in MEG. Inset are scaling factors: multiplying these by 100 (the number of L2/3 pyramidal cells in the model) indicates that 200,000 L2/3 pyramids are sufficient to generate these dipoles.

A scaling factor is applied to the model output (reported in picoampere-meters; pAm) to match units and amplitude in the data (the standard nanampere-meters; nAm), under assumption that the empirical MEG signal is produced by larger populations of synchronous cortical pyramidal neurons than the 200 simulated pyramidal cells. We use this scaling factor to estimate the size of the network that contributes to the recorded response (Jones et al. 2007; Jones et al. 2009; Neymotin et al 2020).

#### Simulating beta event generation and hypothesized recruitment of long-time scale supragranular inhibition

We have previously shown that 50ms bursts of excitatory synaptic input arriving at supragranular layers (i.e. the apical tuft arborization of cortical pyramidal neurons) can explain the prominent negative deflecting peak of the beta event waveshape, see Figure 2Ai (Sherman et al. 2016). Spiking in SI pyramidal cells is not needed to explain these macroscale beta event features, which can arise from the summed subthreshold activity of pyramidal neurons over a large spatial extent. The upstream region creating this burst of input is unspecified in the model. However, a likely source is “higher-order” (cf. “modulatory”, “nonlemniscal”, or “matrix”) thalamus (Mo and Sherman, 2019, Jones, 1998; Jones 2001; Sherman 2016). This hypothesis is bolstered by a line of work showing that nonlemniscal thalamic spikes generate subthreshold effects on both L2/3 and L5b apical dendrites in mice (e.g. Wimmer et al, 2010; Viaene, Petrof and Sherman, 2011; Audette et al, 2017). Importantly, these thalamic inputs concurrently recruit supragranular interneurons, including VIP+ interneurons and potentially other 5HT3a+ interneurons with cell bodies in L2/3 (Audette et al. 2017). The effects on the burst input on the cortical interneuron subnetwork were not accounted for in our previous model (Sherman et al, 2016). Here, we included recruitment of supragranular layer interneurons to examine if and how their recruitment may impact the beta event waveform shape and the relationship between beta events and tactile perception, based on the following motivation.

In brief, the “default” state of cortex is thought to involve tuft inhibition of pyramidal dendrites by apical dendrite-targeting interneurons, e.g. somatostatin-positive (SOM+) interneurons (see Scheyltjens and Arckens, 2016; Figure 5Ai). The immediate higher-order thalamic action on supragranular cortex appears to be disinhibition -- e.g. VIP+ removal of SOM+ inhibition, with calretinin and calbindin playing analogous roles in many primate studies (Meskenaite, 1997; DeFelipe, 1997; Barbas et al, 2018; see also Melchitsky and Lewis, 2008; Krienen et al, 2020) -- and subthreshold pyramidal excitation as mentioned above^†^ (Figure 5Aii). Yet, on longer timescales, strong stimulation of rodent thalamic homolog Pom (see Krubitzer and Kass 1992) profoundly *reduced* sensory-evoked spiking in rodent barrel cortex (Castejon et al. 2016)(Chou et al, 2020), suggesting that the thalamus may act on cortex at multiple timescales by first facilitating and then suppressing activity.

We therefore hypothesized that the thalamic inputs responsible for beta event generation elicit long-timescale inhibition of cortical pyramidal neurons (Figure 5Aiii). Noting that the tactile suppression timescale of ~300ms (Shin et al, 2017) corresponds to that of GABA_B1a_ G-protein coupled inhibition, we furthermore proposed that L2/3 neurogliaform (NGF) cells are one pathway that can mediate this effect (Figure 5Aiii). NGFs are represented among 5HT3a+ interneurons in L2/3 and are coupled to all other interneuron populations via gap junctions (Oláh et al. 2009), Figure 5Aiii. Direct recruitment of NGFs through nonlemniscal bursting is therefore plausible (Audette et al. 2017), but local neurogliaform cells can also be influenced to spike through electrical coupling – thalamic-burst VIP+ recruitment is arguably the single most likely means of activating a gap junction mediated NGF “circuit-breaker” in L2/3. Neurogliaform cells then act perisomatically on L2/3 pyramids, either postsynaptically (Kawaguchi and Kubota 1997; Simon et al. 2005; Overstreet-Wadisch and McBain, 2015; but see also Tremblay, Lee and Rudy 2016) or through volume transmission (Oláh et al. 2009; Overstreet-Wadisch and McBain, 2015; but see also Chittajallu, Pelkey, and McBain, 2013) and on middle-apical dendrites of L5 pyramidal cells (cf. Pérez-Garci et al. 2006). The same effect can be driven by recruitment of those L1 NGFs whose axons ramify mainly in L2/3 (see Figure 3B1 in Wozny and Williams, 2011; cf. Fan, et al 2020).

While the above provides motivation for our model assumptions, we do not assume thalamic bursts are the sole means of causing slow inhibition of the cortical circuit (see Figure 9 and associated discussion), or that NGF cells the only interneuron type that can mediate the slow inhibition. Astrocytes represent another potential source of bulk GABA release that could mediate a GABA_B_ effect, and another candidate for non-GABA_B_ slow inhibition is slow post-inhibitory rebound bursting in SOM+ cells (Audette et al. 2017). *Presynaptic* GABA_B_ time constants are an order of magnitude slower than our behavioral observations (see Pfrieger et al, 1994) and not considered here. The key necessary assumption in the model is that slow inhibition acts on L2/3 pyramids perisomatically and on the mid (or as effectively distal) apical dendrites of the L5 pyramidal neurons, as shown in Figure 5B.

Given the above, we simulated a beta event and its recruitment of slow inhibition as follows. Burst of spikes simultaneously activate excitatory synapses in the SI circuit in a proximal and distal projection pattern, as shown in Figure 5B/Ci and Cii. The proximal drive could emerge from the same higher order thalamus (e.g. see Audette et al. 2017) or from coactivity in a lemniscal thalamic source projecting to granular and infragranular layers. This excitatory drive induces subthreshold currents up and down the pyramidal neuron apical dendrites to generate the beta event waveform shape (Figure Ciii). In addition to subthreshold activation of the pyramidal distal dendrites, the supragranular drive elicits spiking in the L2/3 inhibitory neurons (Figure 5B/C). The inhibitory population is a collapsed representation of interneurons that produce both fast and slow inhibition, e.g. parvalbumin and NGF cells. To represent recruitment of slow inhibition, we simulated a GABA_B_ “synapse” from L2/3 inhibitory cells to the L5 pyramidal neuron middle-apical dendrite in event trials (Figure 5B). All other aspects of the model were identical in event and no-event trials (see Supplementary Table 1 for parameter values and a comparison to earlier model results (Sherman et al. 2016), further results in Supplementary Figures 1 and 2). Figure 5D summarizes the time dependent effects of this beta generation mechanism on a tactile evoked response, as detailed below.

#### Model reproduces the relationship between beta events and tactile evoked responses via multiple latency-dependent effects

Next, we simulated a tactile evoked response during and after a beta event. Our prior work showed that on average events are nearly uniformly distributed across the 1000ms prestimulus period (Shin et al. 2017). We inferred that some events were also likely to occur in the peristimulus interval before and at the time of the arrival of the tactile stimulation to the cortex at 25ms, and therefore modeled beta event centers as occurring between −975ms to 25ms at 100ms increments (t=0 is tactile stimulus onset). We assumed that the macroscale beta events have a dominant impact on the tactile evoked response and for simplicity no inputs were provided to the model other than the thalamic burst drive (Figure 5) and the tactile input sequence (Figure 4). However, similar results were obtained with subthreshold noise inputs to the pyramidal neurons, see Supplementary Figure 3.

There were multiple latency-dependent effects that, when averaged, reproduced the relationship between prestimulus beta events and the MEG SI evoked response (Figure 6A/B). Specifically, the rise after the prominent ~70ms peak was slower and the amplitude of the response around 90ms was smaller for the event trials in both the model and MEG data (compare Figure 6A and B, see also Supplementary Figure 4 for time derivatives). Additionally, the model yielded a lower baseline dipole in the event condition, which is also visible in the MEG data (compare Figure 6A and 6B near 0ms). This initial activity reflects the net downward dipole current occurring during and after a beta event (Figure 5).

**Figure 6.**
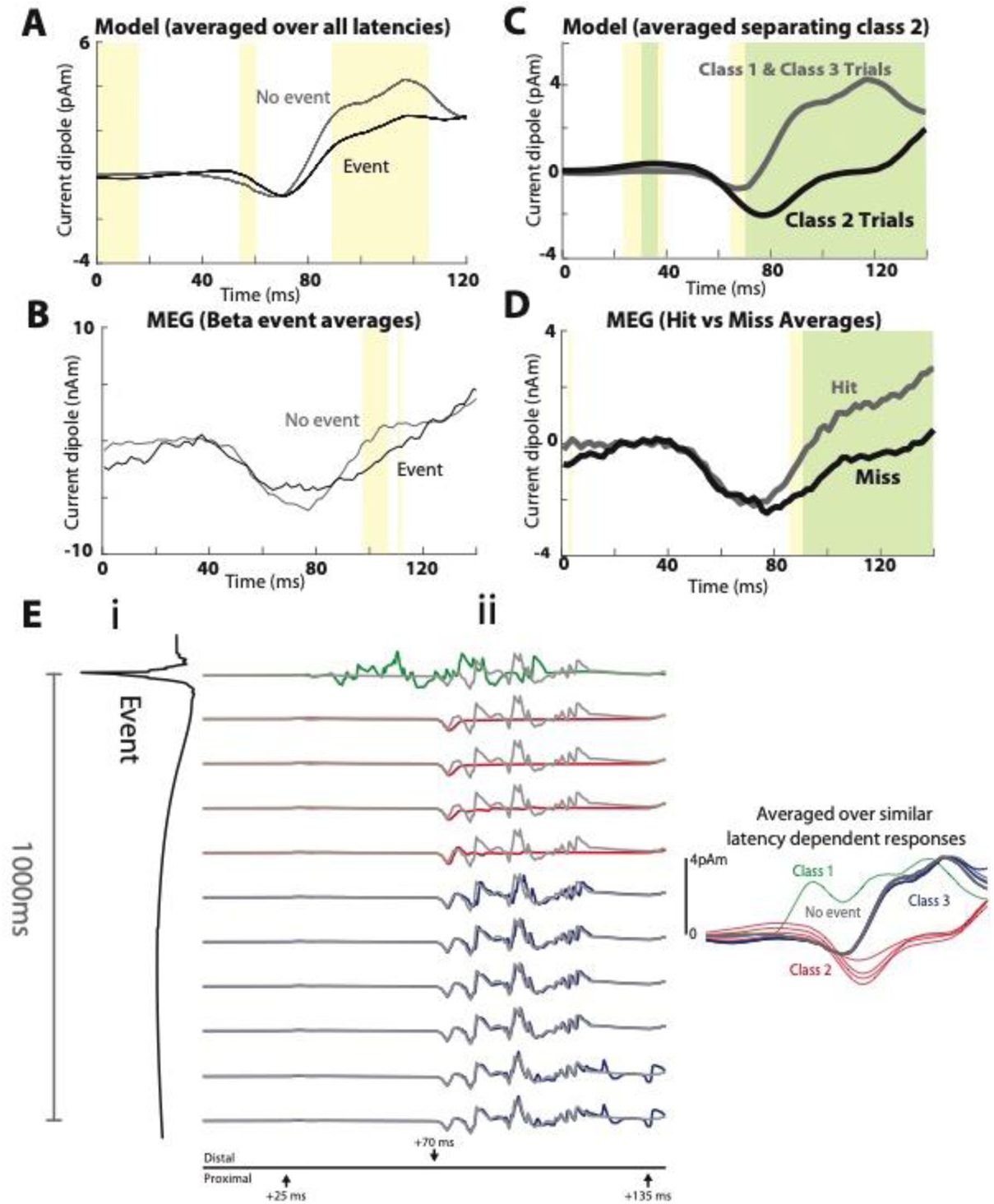
The SI model reproduces the relationship between beta events and tactile evoked responses through three latency-dependent effects. **A)** Model evoked responses averaged over all 11 latencies in (Eii) compared to the evoked response from rest (no event)**. B)** Human MEG SI tactile evoked responses averaged over all trials with or without a beta event in the prestimulus period, as in Figure 3B. **C)** Model evoked responses averaged over class 1 and class 3 trails (assumed hit trials), and over class 2 trials (assumed to be miss trials) **D)** MEG SI tactile evoked responses averaged over detected (hit) and non-detected (miss) trials. **E)** Effects of beta events on model evoked responses depend on latency from the event. **i)** Beta event depicted from event onset to 1000ms after event onset (otherwise identical to Figure 4C). **ii) left:** Raw model evoked responses as a function of latency from event onset. (i) and (ii) are aligned such that (i) is the dipole during and following the event and (ii) is the evoked response when the first sensory input arrives in SI at the timepoint shown in (i). This +25ms shift accounts for conduction delay from the periphery. Time axis in panel ii is the same as in Figure 4A/B. Evoked responses from rest represent no event trials and are shown in grey for comparison. Color-coding represents a by-eye classification of response types. These three latency-dependent patterns are subsequently referred to as Class 1, Class 2 and Class 3 evoked responses. **Right:** Smoothed model evoked responses, corresponding to the 11 raw latency-dependent responses in (Eii). Pointwise significance is shown in yellow (p < 0.05; permutation test) and after FWER-correction in green (α = 0.05; standardized uniform-norm permutation test; see Methods).

A closer look at the latency dependent effects reveals three different evoked response patterns, as color coded in Figure 6Eii: we denote these Class 1 (green), Class 2 (red) or Class 3 (blue); smoothed overlaid responses are shown to the right (see also Supplementary Figure 4). Model Class 1 responses occurred when the thalamic burst and tactile stimulus coincided in SI. Here, the model generated more prominent early dipole activity near 40ms than at other latencies. Class 2 responses occurred when the stimulus arrived between ~50ms-400ms after an event trough. Here, the initial dipole response to the initial feedforward input is nearly flat, with a unitary downward deflection at the time of the 70ms feedback input. Class 3 responses occurred when the stimulus arrived >400ms post-beta-event. These evoked responses were comparable to the no-event responses shown in gray, indicating that the effect of the beta generating mechanisms had worn off after 400ms. Averaging the Class 1 and 3 trials together as model “hit” trials and Class 2 as model “miss” trials reproduced the empirical hit/miss trials without further model tuning (Figure 6C/D); motivation for and further testing of these groupings is detailed in the next Section.

### What are the precise circuit mechanisms underlying the close agreement between the model and data, and how might they causally relate to detection?

We next provide a detailed explanation of the circuit mechanisms generating the model response classes. This examination leads to several new predictions that can be tested with the empirical data. We first examined model spiking activity for each cell type during each class of response and inferred how these responses may correspond to detection correlates in our MEG data. Figure 7A shows spike histograms for each cell type along with raw (lighter) and smoothed (darker) net current dipole during each evoked response class. Layer specific responses are shown in panels B (L2/3) and C (L5). In all panels, spike rates and dipoles are averaged over all latencies for a given class.

**Figure 7:**
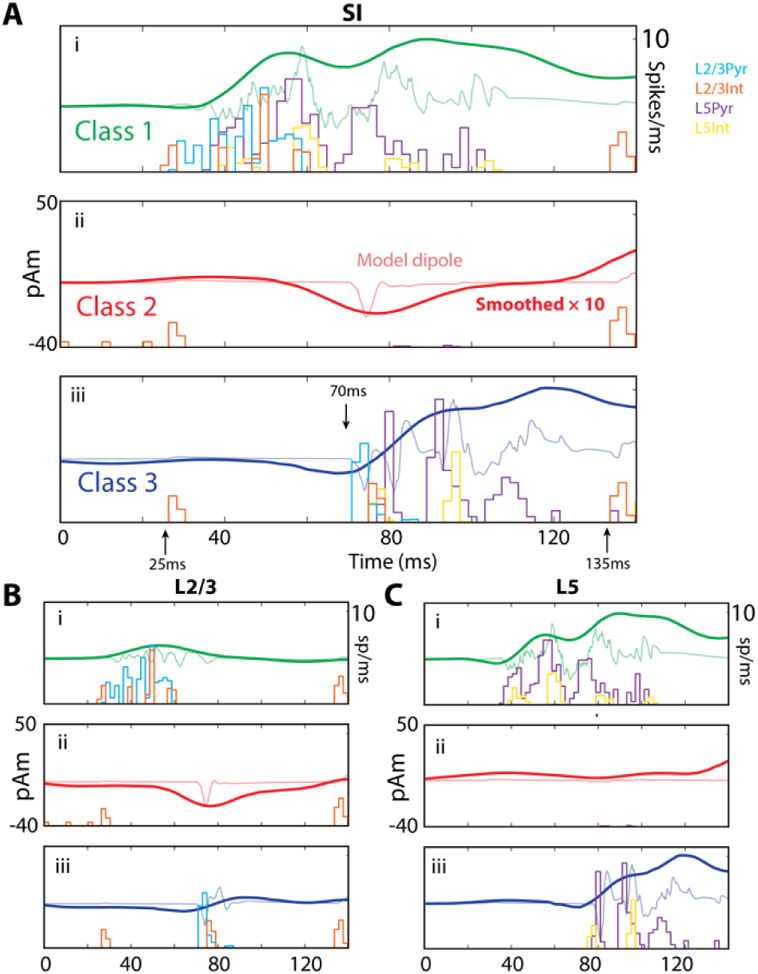
Network spiking activity during three different evoked response classes. **A)** i. Class 1 evoked response averaged over each layer for 101 latencies, using a 10ms rather than the 100ms resolution in Figure 6, and corresponding spike histograms for each cell type in the model. **ii.**and **iii)**. Same as (i), but for Class 2 and Class 3 responses. The Class 1 response shows early and persistent spiking, while the Class 2 response shows suppressed spiking. **B and C)** Contributions to the net current dipole response from L2/3 and L5 separately. **(i)** During the Class 1 response, L2/3 contributes to the early dipole and L5 to the later dipole**. (ii)** During Class 2 responses, L2/3 shows a prominent dipole deflection despite suppression of action potentials in both layers. **(iii)** The Class 3 response is similar to the no beta response in Figure 4C. In each panel, the thin curves show the raw dipole waveform before smoothing and the units of the smooth curve are multiplied by 10 to match the amplitude of the raw dipole.

#### Class 1 responses show early spiking activity and are predicted to be hit trials

During Class 1 responses, feedforward sensory input coincides with depolarization from beta-event excitation of L2/3 pyramidal dendritic tufts and stimulus-driven feedforward inputs on the basal dendrites (Figure 7A-Ci, see also Figures 4/5A). This combined excitation is marked by early spiking activity all L2/3 cells that spreads to spiking activity in all L5 cells, where the spiking persists for a longer period (Figure 7B-Ci). Backpropagating pyramidal action potentials generate the primary positive deflection in the Class 1 dipole response with a high-frequency MEG signature in the unsmoothed signal (green curve Figure 7A).

It is well-known that the excitatory pyramidal neurons in the neocortex relay information to upstream areas including higher order cortical areas, as well as to motor initiation zones, and their activation is critical to somatosensory processing (Vecchia et al, 2020). The high levels of pyramidal spiking activity in the model Class 1 response suggests that tactile information is quickly relayed out of SI to upstream structures, which ultimately allows for registration and report of a stimulus. This finding therefore led us to hypothesize that Class 1 trials correspond to hit trials in our MEG data (Figure 6C/D), and yields a high frequency component that will later aid in validating this prediction with the MEG data.

#### Class 2 responses show a complete abolishment of spiking activity and are predicted to be non-detected (miss) trials

Class 2 trials (Figure 7Aii, Bii, and Cii) were characterized by a near-complete absence of spikes in pyramidal cells of both layers (although isolated doublets from single cells in L5 appeared between 80 and 100ms late in the suppression period). The only spiking response from the initial feedforward input was in L2/3 interneurons — a population that had already been activated by a beta event. Note that the spikes appearing near 0, 10, and 20ms before the ~25ms feedforward input in Figure 7Bii are from the beta-generating thalamic drive. Prestimulus recruitment of these interneurons is responsible for preventing spiking activity in both L2/3 and L5 mainly via the GABA_B_ hyperpolarization.

The observed lack of firing in the pyramidal neurons suggests that, during the Class 2 responses, information about the presence of a tactile stimulus does not exist in SI and thus is not relayed to upstream structures to register and report detection of the stimulus. This finding led us to the hypothesis applied in Figure 6C (and tested further in our MEG data below) that Class 2 trials correspond to miss trials in the MEG data.

#### A closer examination of the mechanisms creating the Class 2 current dipole response

Given that Class 2 responses exhibit a complete lack of pyramidal neuron firing, it is somewhat surprising and non-intuitive that there remains such a strong negative deflection in the dipole response (red curve, Figure 7A). Here, we provide a detailed examination of the layer-specific circuit mechanisms underlying this response. The membrane potentials in each compartment of example L2/3 and L5 pyramidal neurons at ~70ms post-stimulus during a simulated evoked response, without and with a prestimulus beta event, are shown in Figure 8A and 8B respectively. The lack of spiking in L2/3 pyramidal neurons after a beta event was due to perisomatic inhibition evoked by the thalamic-induced spiking in the L2/3 inhibitory neurons (recall Figure 5; compare L2/3 somatic responses in Figure 8Ai/Bi). The lack of spiking in the L5 pyramidal neurons was due to a combination of two processes. One was a lack of excitatory drive from L2/3 to L5 pyramidal neurons, as the L2/3 pyramidal neurons did not spike. The second was the L2/3 inhibition of the L5 pyramidal mid-apical dendrites (see connection in Figure 5A). This inhibition created GABA_B_-mediated attenuation of the L5 response to the 70ms feedback input (compare response in L5 apical dendrites in Figure 8Aii/Bii; note, inhibitory interneuron spiking that provides GABA_B_ input in Figure 8Bii pink shaded regions). In comparison, without a beta event (Figure 8A), L5 pyramidal neurons exhibit a calcium-mediated dendritic burst in response to the 70ms feedback input. This burst starts in the distal dendrites and propagates to the soma (indicated with numbers 1-3 in Figure 8Aii) creating a downward deflecting dipole current near 70ms. We note in this simulation that the calcium-mediated dendritic burst (Figure 8Aii) is only present during the evoked response and not during the beta event generation due to the fact that the conductance of the AMPA drive from the evoked top-down distal input at ~70ms is an order of magnitude larger than the conductance from the thalamic burst generating the beta event, which was specifically tuned so that the response in the pyramidal neurons remains subthreshold (5000pS during evoked distal input, see Figure 4, and 400pS during the beta event, see Figure 5).

**Figure 8:**
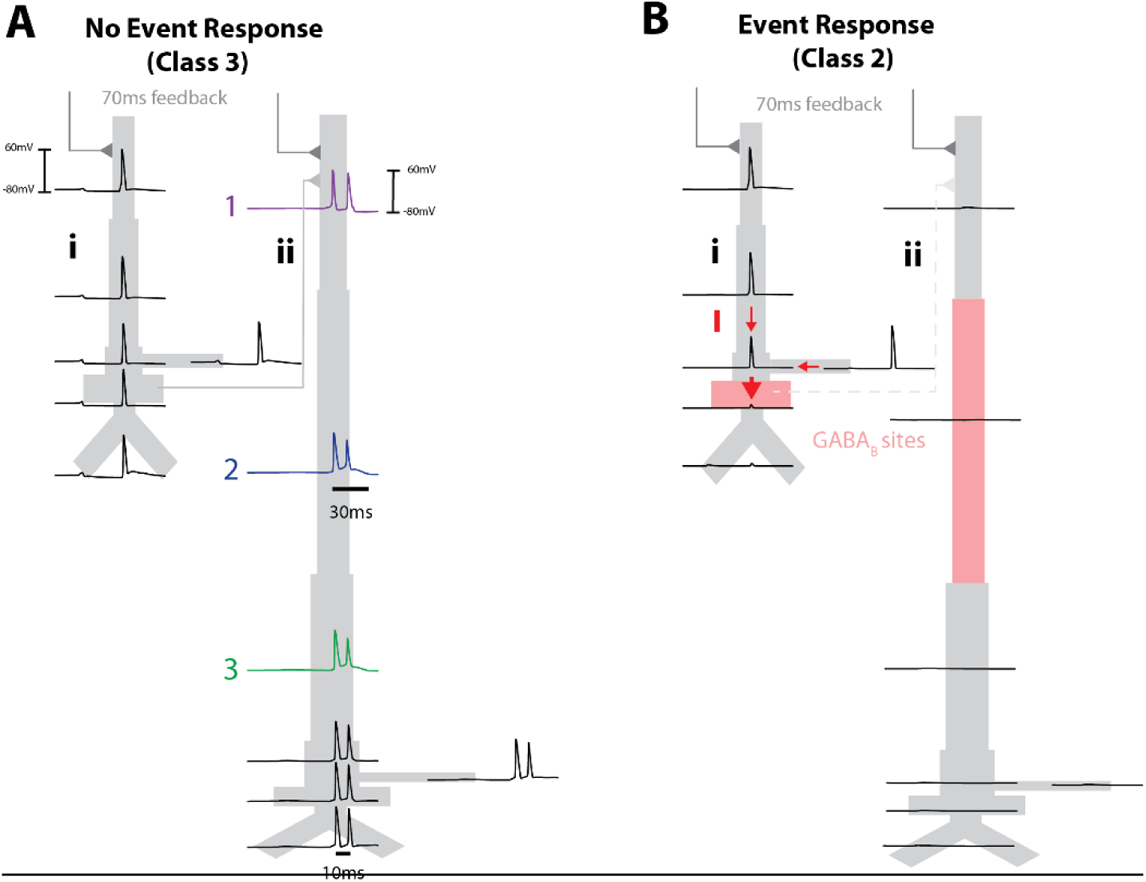
Model Class 2 responses are created by active attenuation of dendritic spikes by strong somatic inhibition. **A)** Membrane potentials are shown for each compartment of exemplar model pyramidal neurons within L 2/3 **(i)** and L5 **(ii)**, responding to stimulus evoked “top-down” feedback arriving 70ms after stimulus onset, without a prestimulus beta event, and with a prestimulus beta event (**B**). Without a beta event, the 70ms feedback input generates dendritic spikes in the pyramidal neurons that propagate to the soma to create negative deflecting dipole currents. Indices 1-3 denote the temporal ordering if of the L5 burst. With a beta event, event-evoked GABA_B1a_ inhibition (Bi) prevents dendritic spikes from reaching the axon initial segment (considered as part of the somatic compartment) in L2/3 pyramids creating large downward dipole currents that generates the Class 2 evoked response shown in Figures 6B and 7, and (Bii) attenuates the feedback response to L5, which prevents any activity in L5 pyramidal neurons. Sites of inhibition are shaded pink; interneurons are not shown. Red arrows in **(Bi)** show that the difference in voltage between the soma and dendritic trunk in L2/3 pyramids creates an effectively maximal dipole.

To understand why the model Class 2 response exhibited a prominent ~70ms deflection without pyramidal neuron firing, we first recall that this peak was created solely by L 2/3 dipole response (Figure 7B, red curves). We examined the difference between L2/3 membrane potentials during no-event and event evoked responses (Figure 8Ai, 8Bi, respectively). Remarkably, the large dipole deflection for Class 2 responses in beta event trials was caused by downward-propagating dendritic spikes in L2/3 pyramidal cells that were actively attenuated at the soma by GABA_B_ suppression; this suppression was not present in the no event case (compare somatic responses in Figure 8Ai/Bi). The voltage difference between the dendritic and somatic compartments creates a large, downward intracellular dipole current in the Class 2 trials (see red arrows Fig 8Bi).

#### Class 3 responses show spiking activity similar to no beta event responses and are predicted to be detected (hit) trials

Class 3 trials (Figure 7Aiii, Biii, Ciii) occurred at latencies where the inhibitory effect during a prestimulus beta event had effectively worn off (i.e. >400ms after the beta event), and waveforms were similar to no-event trials (compare to Figure 4). Details of how the current dipole waveform was generated in this case are described in the description of Figure 4. This finding led us to the hypothesis applied in Figure 6C that Class 3 trails correspond to hit trials in the MEG data.

### Testing two essential model-derived predictions in the MEG data: (i) Class 1 evoked responses correspond to hit trials, and (ii) Class 2 evoked responses correspond to miss trials (Figure 6C/D)

#### Testing Model Prediction i: Class 2 evoked responses correspond to miss trials in the MEG data

To test this prediction, we compared simulated Class 2 evoked responses to averaged evoked responses on miss trials in MEG data. Remarkably, the Class 2 model evoked response was in immediate visual agreement with the averaged evoked response on miss trials with or without a prestimulus beta events (Figure 9). The similarity of the average response on miss trials without an event suggests similar inhibitory mechanisms can and do occur in no-event cases. A linear regression of the model Class 2 response onto the mean response of miss trials revealed similar regression coefficients for each condition and for the average (r^2^=0.74, 0.83, 0.86, respectively; corresponding scaling factors matching the data amplitude are also shown in Figure 9, see detailed methods of correlating Class 2 model waveforms and MEG miss trials in Supplementary Materials and Methods). The model scaling factor is approximately 2000 in all cases; this suggests that approximately 200,000 pyramidal neurons in L2/3 (100 cells in the model multiplied by this factor) are needed to generate the Class 2 response (as L5 did not generate significant dipole activity; see Figure 7Bii).

#### Testing Model Prediction ii: Class 1 evoked responses correspond to hit trials in the MEG data, and are biased to occur during the falling and trough phases of a beta event

We expected Class 1 evoked responses to be rare in our MEG data, as they were not visible in hit trial averages (compare Figure 3B and Figure 7Ai) and because pre-stimulus beta events themselves were observed in only 25% of trials (502 event trials, 1498 no-event trials). However, model details provided several quantifiable targets to find correlates of Class 1 evoked responses (see Figure 10A), and to test the prediction that they are predominantly hit trials.

First, the model predicted that during these Class 1 trials, the first feedforward sensory input to SI (at ~25ms post-stimulus;) coincides with the falling or trough phases of a beta event (Figure 10A; phase interval ~1/4-1/2). For this to be true, beta events would have to reliably occur during the peristimulus period and be aligned such that their troughs occurred near ~25ms post-stimulus on at least a subset of hit trials. We tested this by calculating beta-frequency phase coherence across trials in our MEG data (18-24 Hz see Materials and Methods, Figure 10B). We found that beta-frequency coherence in the post-stimulus period was significantly higher on hit compared to miss trials, with the highest significance precisely at ~25ms post-stimulus (22-32ms, alpha = 0.02; FWER-corrected permutation test). This indicates that a significant fraction of the hit trials had a time locked beta event near the model-predicted timepoints highlighted in Figure 10A. We then verified that the phase of beta frequency activity at 25ms post-stimulus most often occurred near the beta trough on hit trials, namely near phase = ½, by filtering the prestimulus signal in the beta band and generating a histogram of the beta phase at 25ms post-stimulus separately for hit and miss trials (Figure 10B inset).

Second, to find Class 1 responses, we performed a cross correlation analysis between the Materials and Methods). We then examined if those whose onset occurred during ¼ to ½-beta phase (corresponding to the falling or trough phases of a beta event, respectively; see Figure 10A) were hit trials. We found 18 such Class 1 correlates at various phases of a peristimulus beta event in our data (Figure 10C-F). Figure 10C shows an example of the close cross-correlative agreement between the model and one of the identified Class 1 evoked responses in our MEG data. Approximately half of the identified Class 1 responses corresponded to hit trials (10 hit trials and 8 miss trials, green and red, respectively Figure 10F, corresponding phases are as in 10D). However, the hit trial average had a downward sloping dipole during the 0-25ms post stimulus period, while the miss trials did not, consistent with the prediction that hit trials occur during the falling phases of a spontaneous beta event (Figure 10E). Indeed, all hit trials (green Figure 10F) occurred during falling and trough phases of the prestimulus activity (10 of 10). All except for 3 of 8 miss trials (red Figure 10F) occurred outside of falling phases. Miss trials unexplained by the model are shown with pink background in Figure 10F.

Third, during Class 1 evoked responses, the model L2/3 neurons generated a fast oscillation in the average dipole waveform, beginning near ~40ms (see unsmoothed green waveform in Figure 7Ai and 7Bi). The inter-peak interval in the model dipole oscillation was approximately 6ms and high frequency coherence across trials (160-200Hz) occurred with a peak near ~40ms (Figure 10G). This led to the prediction that Class 1 correlated hit trials in the MEG data should have a reliable (i.e. stimulus-locked) fast oscillating response in the dipole waveform with high-frequency cross-trial coherence that is not present in miss trials. Indeed, we found higher 160-200Hz cross-trial coherence on hit compared to miss trials, and miss trial coherences were nearly zero throughout the early evoked response period (Figure 10H; alpha = 0.05; FWER-corrected permutation test). The timing of the significant differences between hit and miss trials in the MEG data peaked near 40ms lOG and lOH).

## Discussion

Building from prior work showing prestimulus SI beta events predict nondetection for ~300-400ms after an event (Shin et al. 2017), we set out to understand the cellular and circuit level mechanisms by which this sensory suppression may occur. Biophysically principled neural modeling combined with human MEG offered a precise explanation: Higher-order thalamic bursts that generate beta events also indirectly activate supragranular pyramidal GABA_B1a_ receptors. This suppresses spiking throughout the cortical circuit, inhibiting sensory perception after the beta event for the timescale of GABA_B1a_ effects, ~400ms.

The model also showed that when a “top-down” beta-generating thalamic burst and a tactile stimulus arrive in SI at the same time (a *coincident* or *on-stimulus* beta event), the coincidence generates an early cascade of L2/3 spiking, followed by L5 pyramidal spiking. This cascade created predicted MEG features that were verified as correlates of perceived (hit) trials by three separate means of testing (Figure 10). This result requires a reinterpretation of beta’s role beyond pure inhibition, as it indicates a mechanism by which the thalamocortical activity underlying beta events can facilitate perception when timed appropriately.

### Consistency of the current findings with prior studies

It is well known that features of sensory-evoked responses can correlate with successful perception, and that spontaneous dynamic brain states, often described in terms of prestimulus “oscillations”, influence this correlation (Palva et al. 2005; Jones et al. 2007; Jones et al. 2009). However, an understanding of the precise neural mechanisms that could provide a causal link between prestimulus brain states and perception is lacking, particularly in humans where invasive recordings are rare. Our modeling framework enabled interpretation of cellular and circuit-level mechanisms underlying the effect of beta-event processes on tactile evoked response and perception in humans via direct comparison of model and source-localized MEG data, in identical physical units. We argue for a causal relationship between these transient beta-event processes (i.e. cortical excitation and inhibition induced by thalamic bursts) and tactile detection, by providing a detailed model dissection of processes underlying the observed neural dynamics.

A critical model extension here was based on the novel hypothesis that higher-order thalamic bursts recruit slow GABA_B1a_-mediated inhibition in the post-event, prestimulus time period. Updating our prior model to reflect this hypothesis was crucial to our conclusions regarding thalamic burst-mediated tactile suppression and did not significantly change the robustly observed stereotypical beta event waveform shape (Sherman et al. 2016; Little et al. 2018; but see also Supplementary Figures 1 and 2). Our findings are also in agreement with several studies that have suggested slow inhibition in the supragranular layers — including through GABA_B_ mechanisms — is a key regulator of conscious perception (Craig and McBain 2014; Cone et al. 2019) and is specifically involved in inhibiting somatosensory perception (Pluta et al. 2019). For example, Larkum and colleagues have shown that dendrite-mediated suppression of rodent somatosensory perception is regulated by GABA_B_ receptor activation in the mid-apical dendrite of L5 pyramidal cells, driven by a transcallosal inhibitory projection (Larkum et al. 1999; Takahashi et al. 2016). Our results extend theirs by modeling a local circuit mechanism for this inhibition in humans, noteworthy because long-range inhibitory projections are prevalent in rodents but have no known analog in primates (Barbas et al. 2018).

The agreement between model and MEG data support a causal influence of the processes underlying transient beta events on evoked correlates of perception, but it does not account for other post-stimulus dynamics that may be independent of prestimulus beta events. For instance, in a previous study (Jones et al. 2007), we showed that decreasing the arrival time and increasing the strength of the post-stimulus ~70ms distal “feedback” and subsequent ~135ms “re-entrant” thalamic inputs to SI (Figure 4) could also account for evoke correlates of perception, without mechanisms alone could generate the observed differences in the evoked response, but we do not rule out the possibility that a combination of pre-stimulus and post-stimulus influences contribute to the results.

### Anticipatory beta events as a predictive signaling and learning mechanism

Our results suggest thalamic bursts generating SI beta events can both inhibit and enhance threshold-level tactile perception in a corresponding somatotopic region. Specifically, our findings suggest that if a beta event coincides precisely with tactile information, perception is enhanced, but after this time window closes, information processing is suppressed by slow inhibitory currents. Although on-stimulus beta events were rare in the MEG data, this dichotomy raises an important conceptual debate: Are beta events (viz. thalamic bursts) actively engaged with task demands to suppress perception? Or, does beta reflect a limited time window for *enhancement* of perception, with sensory suppression a signature of mistimed (too-early) beta events? Is there an advantage to brief enhancement of sensory stimuli followed by prolonged inhibition? Our study was not designed to definitively address the cognitive strategy by which the beta process is temporally engaged, but we shall discuss interpretations of beta in timing prediction, and suggest that regardless of the temporal strategy by which beta is engaged, one role of the beta process may be to link detection with the learning of timing.

Many studies have shown that beta (and therefore its underlying neural process) can be deployed with temporal specificity in coordination with task and cognitive demands (van Ede et al. 2011; Fujioka et al. 2012; Sacchet et al. 2015; Spitzer and Haegens 2017; Fiebelkorn et al. 2018; Little et al. 2018). In the sensorimotor domain, we and others have observed that beta power decreases in localized somatotopic brain regions during the anticipatory period following a cue to attend to a corresponding body location, and increases in somatotopic regions corresponding to non-attended body locations (Linkenkaer-Hansen et al. 2004; Jones et al. 2010; van Ede et al. 2011). Such beta power modulation is also synchronized with frontal cortex (Sacchet et al. 2015), and peripheral muscles with or without task-specific motor demands (van Ede and Maris 2013). Lower averaged prestimulus/anticipatory beta power is associated with faster responses (van Ede et al. 2011), and higher detection rates (Jones et al. 2010; Shin et al. 2017), phenomena that have also been observed in visual-motion detection tasks where beta appears to track evidence accumulation (Donner et al. 2009). Moreover, it has been shown that beta power is actively decreased in a temporally specific manner, when a tactile stimulus is expected but not delivered (van Ede et al. 2011). These and other studies (see also studies of motor cortex beta oscillations, where decreased beta activity corresponds with more certain responses and faster reaction times (Little et al. 2018)) suggest sensorimotor beta is inhibitory to function and actively decreased to enhance somatosensory detection, and/or actively increased to block irrelevant information.

The above results are compatible with the view that top-down processes underlying beta event generated are recruited with temporal precision for attentional suppression of irrelevant information, perhaps mediated by the effects of superior colliculus on higher-order thalamic nuclei (Gharaei et al, 2020). In this case, the faciliatory Class 1 responses arising from stimulus/event coincidence could be interpreted as an “error” signal indicating that inhibition arrived too late, and that the beta event timing should be adjusted. Alternatively, the brain may be trying to engage thalamic bursts at the “predicted” time of the stimulus in order to facilitate perception of weak stimuli, in which case prestimulus beta events represent premature timing predictions and the suppressed Class 2 response could be interpreted as the “error” signal. It is worth noting here that the aforementioned studies showing an inverse relation between beta power and perceptibility/motor action do not report beta phase. It is possible that smaller-amplitude events, with the appropriate phase alignment, are sufficient to facilitate processing when a prediction can be precisely made.

In either case, according to our model, on-stimulus detected beta events represent a coincidence of “top-down” subthreshold excitatory input to apical dendrites and “bottom-up” drive to basal dendrites in L2/3 pyramidal neurons, aligning with a matching principle posed as a necessary component of conscious perception (Grossberg 1980). Our results suggest an explicit mechanism for this abstract principle, specifying that a dendritic coincident match should be followed by a time-locked increase in spiking activity in L2/3, which later initiates L5 bursts, which themselves have been argued to link a percept globally across the cortex (Takahashi et al. 2016; see also Aru, Suzuki and Larkum, 2020). Indeed, L2/3 to L5 propagation was recently directly observed in a visual discrimination task, which suggests this process may be a global requisite for perception (Marshel et al, 2019; but see also Takahashi et al, 2020).

Theoretical considerations based in part on our model results further suggest that such a temporal matching mechanism could appear in concert with a learning mechanism that processes timing “errors” — supporting a role beyond pure sensory suppression for inhibition recruited in the post-event period. Specifically, recall that during suppressed Class 2 responses remarkably large dipole currents remain, due to layer 2/3 dendritic spikes that propagate toward the soma but are extinguished by perisomatic inhibition (Figure 8), consistent with prior findings suggesting inhibition can generate large field signals (Telenczuk et al, 2017). Several lines of evidence suggest that this process may provide a substrate for engaging “one-shot” postsynaptic learning, as discussed further in the next section, in line with recent work indicating a causal relationship between beta events and plasticity (Zanos et al. 2018). Either or both of the facilitated (Class 1) or suppressive (Class 2) phases could be associated with learning mechanisms, although we shall focus on a putative mechanism for learning from the Class 2 response in the section to follow.

Further studies are needed to show if and how the beta-event process is actively modulated to suppress, amplify, and/or to learn the timing of sensory and/or internally-generated signals. Our present work extends and refines a variety of previous studies showing that this process is dynamically engaged to meet task demands. The upstream sources responsible for beta events — in particular higher-order thalamic bursts — can modulate cortical activity to gate the perceptual process at threshold and help regulate learning. The work here points to new and precise cortical circuit mechanisms that can mediate this gating.

### Mechanisms of learning during the inhibitory phase of the beta-event process

A natural pathway for thalamic bursts to induce postsynaptic learning as proposed above is through the VIP+ interneuron system, known to be activated by higher-order thalamus (Audette et al. 2017) and recently shown to mediate NMDA-based long term potentiation (LTP) in pyramidal cells (Williams and Holtmaat 2018). Though not explicitly modeled, we hypothesized that the aforementioned VIP networks, when synchronized, also recruit neurogliaform cells via electrical synapses, which could cause beta event suppression in our model (see discussion in results and Figure 5). In rodent somatosensory L2/3 pyramidal cells, NMDA spike-mediated LTP, also with a thalamic source, had been found in the *absence* of somatic spiking activity (Gambino et al. 2014). This parallels our model (which does not include long-term plasticity), where L2/3 inhibition with a thalamic source results in a large dipole signal in the absence of somatic spiking (Figure 8).

Along these lines, it is interesting to note that increased somatic calcium flow has been reported under strong somatic inhibition during slow-wave sleep, where it was hypothesized to optimize learning (Niethard et al. 2018). In that study, GABA_A_-ergic parvalbumin positive interneurons were found responsible for the effect – however, GABA_B_ activation tends to follow hyperactivation of GABA_A_ interneurons as GABA overflows the synaptic cleft (Scanziani 2000).

Therefore, in addition to this VIP-NMDA mechanism, it is possible that the collision between dendritic spikes and perisomatic suppression may lead to calcium transfer to the soma (cf. Chiu et al, 2020); such signaling cation fluxes are well-known to play fundamental and general roles in structural plasticity (Berridge 1998; Augustine et al. 2003).

In summary, together with our modeling results, the literature appears to support two beta-event associated learning mechanisms — one short-term process via VIP - NMDA at the dendrite, and one long-term process via calcium at the soma — both of which might be linked through the Class 2 mechanism. It is possible that VIP-coupled NMDA may act as a short-term tag on a synaptic site, which could later be converted to permanent memory after dendritic spike suppression and subsequent somatic calcium influx, mediated by neurogliaform cells and/or soma-targeting GABA_B_-ergic populations.

### GABA_B_ may govern beta event durations and inter event intervals

Beta-associated recruitment of GABA_B_ inhibition may be used to set the lifetimes of beta events themselves. That is, if beta events involve a recurrent circuit between cortex and thalamus, the timescale of recurrent activation in that circuit may be limited by GABA_B_ activation. Remarkably, this timescale accords approximately with both the mean event duration (~150ms latency to peak GABA_B_ conductance vs. ~150ms event duration) and inter-event intervals (~250ms; approximately the GABA_B_ fall time constant) reported in (Shin, et al, 2017). Further, extended recurrent oscillations in the beta band -- in L2/3 -- were found to be caused by stimulation of rodent thalamic homolog POm (see Figure 5 in Zhang and Bruno, 2019). These extended oscillations were discovered under anesthesia with fentanyl, an opioid. As opioids cause NGF populations to cease firing (Krook-Magnusen, et al 2011), their use could turn off the suppressive effect of beta events yielding longer thalamocortical recurrences, and hence repeated beta-events under our framework.

### Modeling Assumptions, Limitations and Independence

This study builds from a body of prior MEG and modeling work, where we first showed that post-stimulus features of the tactile-evoked response in SI (i.e. the M70 amplitude and slope) alone could, in principle, account for correlates of tactile detection without considering prestimulus state (Jones et al. 2007). Later, we established that prestimulus low-frequency rhythms (i.e. the SI mu rhythm, comprised of 7-14 Hz alpha and 15-29 Hz beta rhythms) influence components of the evoked response through specific network mechanisms, including a strong inhibitory influence mediated by sensory evoked inhibition (Jones et al. 2009). However, in the latter study, we reported only on averaged data, and did not separate the effects of the alpha and beta components of the SI mu rhythm, nor did we investigate the relation of these effects to perception. Further studies showed alpha and beta have separable effects on perception and attention (Jones et al. 2010; Sacchet et al. 2015). The current study is the first to look at circuit mechanism mediated perceptual effects specifically in the beta band.

The chosen SI model configuration is grounded in generalizable principles of cortical circuitry and known somatosensory cortical architecture. Some of the model assumptions create limitations in our conclusions, while many of the findings are independent of specific model choices. One potential limitation is that we simulate only one type of GABA_B_ receptor. We found that even at low densities, simulated GABA_B1a_ channels in middle-apical dendrites of L5 pyramidal cells induced by beta-generating mechanisms can prevent these cells from firing during sensory stimulation. However, the primary target of L2/3 NGF activation on L5b apical dendrites is presumably the GABA_B1b_ receptor, which inactivates calcium channels while admitting sodium spike propagation (Pérez-Garci et al. 2006). As such, it is possible that L5 pyramidal spikes can be recruited by weak sensory stimuli in the absence of L2/3 recruitment when they are close to their firing threshold. Our model, and the assumed higher-order thalamic origin of burst events, also does not account for the higher-order thalamic recruitment of L5a pyramidal spikes observed in rodent slices (Audette et al. 2017). Despite these assumptions, it is crucial to note that one of our main results, while dependent on our proposed beta-generating mechanism – namely, the ~50ms burst of subthreshold excitatory synaptic input to pyramidal neuron distal dendrites -- are essentially independent of free parameters in the model. That is, the non-detected Class 2 waveforms should occur in *any* model that contains dendritic geometry, dendritic spikes, and strong perisomatic inhibition.

We have assumed beta events are mediated by higher-order thalamus, but it is possible that “top-down” corticocortical connections play a role in generating beta events as well. The source of the distal (cf. modulatory) input does not change the fundamental findings of our study, which identifies neural circuit mechanisms generating beta-mediated evoked response correlates of perception within a canonical cortical unit. Finally, while there is close agreement between the model results and the MEG data, all model results are still “predictions” on the circuit dynamics generating that data. Validation of these predictions requires both out-of-sample confirmation and/or testing with invasive recordings (e.g, laminar recordings as in Sherman et al 2016), or with other imaging modalities (e.g. laminar MEG, fMRI spectroscopy, tractography, etc). Our modeling framework provides targeted network features to guide future research on the role of beta rhythms, and their underlying sources, in sensory perception.

## Tables

See Supplementary Materials and Methods, Supplementary Table 1

## Acknowledgements

We thank Professor Matt Harrison for a number of helpful discussions and explanations, and for providing MATLAB code for the permutation and modified max-T tests. We thank Chris Black for assistance with Figure 1.

## Footnotes

* Here, we use “event” rather than “burst” so as not to confuse events with their mechanistic generators, which we believe to be bursts in upstream sources. Our interpretation of events is as “sequences of bursts”, viz. “bursts of bursts” where the length of the sequence can be as low as one.

† Of these, only the excitation will substantially affect the dipole as the resting (disinhibitory) potential is near the chloride reversal (GABAA inhibitory) potential. We therefore do not explicitly model the SOM+/VIP+ interaction.

## References

Amarasingham A, Harrison MT, Hatsopoulos NG, Geman S. 2011 Nov 18. Conditional Modeling and the Jitter Method of Spike Re-sampling: Supplement.

Audette NJ, Urban-Ciecko J, Matsushita M, Barth AL. 2017. POm Thalamocortical Input Drives Layer-Specific Microcircuits in Somatosensory Cortex. Cereb Cortex.:1–17.

Augustine GJ, Santamaria F, Tanaka K. 2003. Local Calcium Signaling in Neurons. Neuron. 40(2):331–346. doi:10.1016/S0896-6273(03)00639-1.

Barbas H, Wang J, Joyce MKP, García-Cabezas MÁ. 2018. Pathway mechanism for excitatory and inhibitory control in working memory. J Neurophysiol. 120(5):2659–2678. doi:10.1152/jn.00936.2017.

Berridge MJ. 1998. Neuronal calcium signaling. Neuron. 21(1):13–26.

Carracedo, L. M., Kjeldsen, H., Cunnington, L., Jenkins, A., Schofield, I., Cunningham, M. O., … & Whittington, M. A. (2013). A neocortical delta rhythm facilitates reciprocal interlaminar interactions via nested theta rhythms. Journal of Neuroscience, 33(26), 10750–10761.

Castejon C, Barros-Zulaica N, Nuñez A. 2016. Control of somatosensory cortical processing by thalamic posterior medial nucleus: a new role of thalamus in cortical function. PLoS One. 11(1):e0148169.

Cauller LJ, Kulics AT. 1991. The neural basis of the behaviorally relevant N1 component of the somatosensory-evoked potential in SI cortex of awake monkeys: evidence that backward cortical projections signal conscious touch sensation. Exp brain Res. 84(3):607–619.

Chen D, Fetz EE. 2005. Characteristic Membrane Potential Trajectories in Primate Sensorimotor Cortex Neurons Recorded In Vivo. J Neurophysiol. 94(4):2713–2725. doi:10.1152/jn.00024.2005.

Chou, X. L., Fang, Q., Yan, L., Zhong, W., Peng, B., Li, H., … & Zhang, L. I. (2020). Contextual and cross-modality modulation of auditory cortical processing through pulvinar mediated suppression. Elife, 9, e54157.

Cole SR, van der Meij R, Peterson EJ, de Hemptinne C, Starr PA, Voytek B. 2017. Nonsinusoidal Beta Oscillations Reflect Cortical Pathophysiology in Parkinson’s Disease. J Neurosci. 37(18):4830–4840. doi:10.1523/JNEUROSCI.2208-16.2017.

Cole SR, Voytek B. 2017. Brain Oscillations and the Importance of Waveform Shape. Trends Cogn Sci. 21(2):137–149. doi:10.1016/j.tics.2016.12.008.

Cone JJ, Scantlen MD, Histed MH, Maunsell JHR. 2019. Different Inhibitory Interneuron Cell Classes Make Distinct Contributions to Visual Contrast Perception. eNeuro. 6(1):1–12. doi:10.1523/ENEURO.0337-18.2019.

Craig MT, McBain CJ. 2014. The emerging role of GABAB receptors as regulators of network dynamics: fast actions from a ‘slow’ receptor? Curr Opin Neurobiol. 26:15–21.

DeFelipe, J. (1997). Types of neurons, synaptic connections and chemical characteristics of cells immunoreactive for calbindin-D28K, parvalbumin and calretinin in the neocortex. Journal of chemical neuroanatomy, 14(1), 1–19.

Donner TH, Siegel M, Fries P, Engel AK. 2009. Buildup of choice-predictive activity in human motor cortex during perceptual decision making. Curr Biol. 19(18):1581–1585. doi:10.1016/j.cub.2009.07.066.

van Ede F, de Lange F, Jensen O, Maris E. 2011. Orienting attention to an upcoming tactile event involves a spatially and temporally specific modulation of sensorimotor alpha-and beta-band oscillations. J Neurosci. 31(6):2016–2024.

van Ede F, Maris E. 2013. Somatosensory Demands Modulate Muscular Beta Oscillations, Independent of Motor Demands. J Neurosci. 33(26):10849–10857. doi:10.1523/JNEUROSCI.5629-12.2013.

van Ede F, Quinn AJ, Woolrich MW, Nobre AC. 2018. Neural Oscillations: Sustained Rhythms or Transient Burst-Events? Trends Neurosci. 41(7):415–417. doi:10.1016/j.tins.2018.04.004.

Engel AK, Fries P. 2010. Beta-band oscillations — signalling the status quo? Curr Opin Neurobiol. 20(2):156–165. doi:10.1016/J.CONB.2010.02.015.

Fan, L. Z., Kheifets, S., Böhm, U. L., Wu, H., Piatkevich, K. D., Xie, M. E., … & Cohen, A. E. (2020). All-optical electrophysiology reveals the role of lateral inhibition in sensory processing in cortical layer 1. Cell, 180(3), 521–535.

Feingold J, Gibson DJ, DePasquale B, Graybiel AM. 2015. Bursts of beta oscillation differentiate postperformance activity in the striatum and motor cortex of monkeys performing movement tasks. Proc Natl Acad Sci. 112(44):13687–13692. doi:10.1073/pnas.1517629112.

Fiebelkorn IC, Pinsk MA, Kastner S. 2018. A Dynamic Interplay within the Frontoparietal Network Underlies Rhythmic Spatial Attention. Neuron. 99(4):842–853. doi:https://doi.org/10.1016/j.neuron.2018.07.038.

Fujioka T, Trainor LJ, Large EW, Ross B. 2012. Internalized timing of isochronous sounds is represented in neuromagnetic β oscillations. J Neurosci. 32(5):1791–1802. doi:10.1523/JNEUROSCI.4107-11.2012.

Gambino F, Pagès S, Kehayas V, Baptista D, Tatti R, Carleton A, Holtmaat A. 2014. Sensory-evoked LTP driven by dendritic plateau potentials in vivo. Nature. 515:116–119.

De Gennaro L, Ferrara M. 2003. Sleep spindles: an overview. Sleep Med Rev. 7(5):423–440.

Fries, Pascal. “Rhythms for cognition: communication through coherence.” Neuron 88.1 (2015): 220–235.

Gharaei, S., Honnuraiah, S., Arabzadeh, E., & Stuart, G. J. (2020). Superior colliculus modulates cortical coding of somatosensory information. Nature communications, 11(1), 1–14. doi:10.1038/s41467-020-15443-1

Grossberg S. 1980. How does a brain build a cognitive code? Psychol Rev. 87(1):1–51. doi:10.1037/0033-295X.87.1.1.

Iemi, L., Busch, N. A., Laudini, A., Haegens, S., Samaha, J., Villringer, A., & Nikulin, V. V. (2019). Multiple mechanisms link prestimulus neural oscillations to sensory responses. Elife, 8, e43620.

Jenkinson N, Brown P. 2011. New insights into the relationship between dopamine, beta oscillations and motor function. Trends Neurosci. 34(12):611–618. doi:10.1016/j.tins.2011.09.003.

Jestrović I, Coyle JL, Sejdić E. 2014. The effects of increased fluid viscosity on stationary characteristics of EEG signal in healthy adults. Brain Res. 1589:45–53. doi:10.1016/j.brainres.2014.09.035.

Jiang X, Shen S, Cadwell CR, Berens P, Sinz F, Ecker AS, Patel S, Tolias AS. 2015. Principles of connectivity among morphologically defined cell types in adult neocortex. Science (80-). 350(6264):aac9462. doi:10.1126/science.aac9462.

Jones EG. 2001. The thalamic matrix and thalamocortical synchrony. Trends Neurosci. 24(10):595–601. doi:10.1016/S0166-2236(00)01922-6.

Jones SR. 2011. Biophysically Principled Computational Neural Network Modeling of Magneto-/Electro-Encephalography Measured Human Brain Oscillations. In: Neuronal Network Analysis. Springer. p. 459–485.

Jones SR. 2016. When brain rhythms aren’t ‘rhythmic’: implication for their mechanisms and meaning. Curr Opin Neurobiol. 40:72–80. doi:10.1016/j.conb.2016.06.010.

Jones SR, Kerr CE, Wan Q, Pritchett DL, Hämäläinen M, Moore CI. 2010. Cued spatial attention drives functionally relevant modulation of the mu rhythm in primary somatosensory cortex. J Neurosci. 30(41):13760–13765.

Jones SR, Pritchett DL, Sikora MA, Stufflebeam SM, Hämäläinen M, Moore CI. 2009. Quantitative Analysis and Biophysically Realistic Neural Modeling of the MEG Mu Rhythm: Rhythmogenesis and Modulation of Sensory-Evoked Responses. J Neurophysiol. 102(6):3554–3572. doi:10.1152/jn.00535.2009.

Jones SR, Pritchett DL, Stufflebeam SM, Hämäläinen M, Moore CI. 2007. Neural correlates of tactile detection: a combined magnetoencephalography and biophysically based computational modeling study. J Neurosci. 27(40):10751–10764.

Kawaguchi Y, Kubota Y. 1997. GABAergic cell subtypes and their synaptic connections in rat frontal cortex. Cereb Cortex. 7(6):476–486. doi:10.1093/cercor/7.6.476.

de Kock CPJ, Sakmann B. 2008. High frequency action potential bursts (≥ 100 Hz) in L2/3 and L5B thick tufted neurons in anaesthetized and awake rat primary somatosensory cortex. J Physiol. 586(14):3353–3364. doi:10.1113/jphysiol.2008.155580.

Krook-Magnuson, E., Luu, L., Lee, S. H., Varga, C., & Soltesz, I. (2011). Ivy and neurogliaform interneurons are a major target of μ-opioid receptor modulation. Journal of Neuroscience, 31(42), 14861–14870.

Krienen, F. M., Goldman, M., Zhang, Q., Del Rosario, R. C., Florio, M., Machold, R., … & McCarroll, S. A. (2020). Innovations present in the primate interneuron repertoire. Nature, 586(7828), 262–269.

Krubitzer, L. A., & Kaas, J. H. (1992). The somatosensory thalamus of monkeys: cortical connections and a redefinition of nuclei in marmosets. Journal of Comparative Neurology, 319(1), 123–140.

Larkum ME, Zhu JJ, Sakmann B. 1999. A new cellular mechanism for coupling inputs arriving at different cortical layers. Nature. 398(6725):338–341. doi:10.1038/18686.

Lehmann EL, Romano JP. 2006. Testing statistical hypotheses. Springer Science & Business Media.

Linkenkaer-Hansen K, Nikulin V V., Palva S, Ilmoniemi RJ, Palva JM. 2004. Prestimulus oscillations enhance psychophysical performance in humans. J Neurosci. 24(45):10186–10190.

Little S, Bonaiuto J, Barnes G, Bestmann S. 2018. Motor cortical beta transients delay movement initiation and track errors. bioRxiv. doi:10.1101/384370.

Lundqvist M, Herman P, Warden MR, Brincat SL, Miller EK. 2018. Gamma and beta bursts during working memory readout suggest roles in its volitional control. Nat Commun. 9(1). doi:10.1038/s41467-017-02791-8.

Lundqvist M, Rose J, Herman P, Brincat SLL, Buschman TJJ, Miller EKK. 2016. Gamma and Beta Bursts Underlie Working Memory. Neuron. 90(1):152–164. doi:10.1016/j.neuron.2016.02.028.

Marshel, J. H., Kim, Y. S., Machado, T. A., Quirin, S., Benson, B., Kadmon, J., … & Deisseroth, K. (2019). Cortical layer–specific critical dynamics triggering perception. Science, 365(6453), eaaw5202.

Melchitzky, D. S., & Lewis, D. A. (2008). Dendritic-targeting GABA neurons in monkey prefrontal cortex: Comparison of somatostatin-and calretinin-immunoreactive axon terminals. Synapse, 62(6), 456–465.

Meskenaite, V. (1997). Calretinin-immunoreactive local circuit neurons in area 17 of the cynomolgus monkey, Macaca fascicularis. Journal of Comparative Neurology, 379(1), 113–132.

Mo, C., & Sherman, S. M. (2019). A sensorimotor pathway via higher-order thalamus. Journal of Neuroscience, 39(4), 692–704.

Moreau D, Lefort C, Pas J, Bardet SM, Leveque P, O’Connor RP. 2018. Infrared neural stimulation induces intracellular Ca2+ release mediated by phospholipase C. J Biophotonics. 11(2):e201700020. doi:10.1002/jbio.201700020.

Neymotin, S. A., Daniels, D. S., Caldwell, B., McDougal, R. A., Carnevale, N. T., Jas, M., … & Jones, S. R. (2020). Human Neocortical Neurosolver (HNN), a new software tool for interpreting the cellular and network origin of human MEG/EEG data. Elife, 9, e51214. doi:10.7554/eLife.51215

Nichols T, Hayasaka S. 2003. Controlling the familywise error rate in functional neuroimaging: a comparative review. Stat Methods Med Res. 12(5):419–446. doi:10.1191/0962280203sm341ra.

Niethard N, Ngo H-V V., Ehrlich I, Born J. 2018. Cortical circuit activity underlying sleep slow oscillations and spindles. Proc Natl Acad Sci. 115(39):E9220–E9229. doi:10.1073/pnas.1805517115.

Oláh S, Füle M, Komlósi G, Varga C, Báldi R, Barzó P, Tamás G. 2009. Regulation of cortical microcircuits by unitary GABA-mediated volume transmission. Nature. 461(7268):1278–1281.

Otis TS, De Koninck Y, Mody I. 1993. Characterization of synaptically elicited GABAB responses using patch-clamp recordings in rat hippocampal slices. J Physiol. 463(1):391–407.

Palva S, Linkenkaer-Hansen K, Näätänen R, Palva JM. 2005. Early neural correlates of conscious somatosensory perception. J Neurosci. 25(21):5248–5258.

Pérez-Garci E, Gassmann M, Bettler B, Larkum ME. 2006. The GABAB1b isoform mediates long-lasting inhibition of dendritic Ca2+ spikes in layer 5 somatosensory pyramidal neurons. Neuron. 50(4):603–616.

Pfrieger, F. W., Gottmann, K., & Lux, H. D. (1994). Kinetics of GABAB receptor-mediated inhibition of calcium currents and excitatory synaptic transmission in hippocampal neurons in vitro. Neuron, 12(1), 97–107.

Pluta SR, Telian GI, Naka A, Adesnik H. 2019. Superficial Layers Suppress the Deep Layers to Fine-tune Cortical Coding. J Neurosci. 39(11):2052–2064. doi:10.1523/JNEUROSCI.1459-18.2018.

Rule ME, Vargas-Irwin CE, Donoghue JP, Truccolo W. 2017. Dissociation between sustained single-neuron spiking and transient β-LFP oscillations in primate motor cortex. J Neurophysiol. 117(4):1524–1543. doi:10.1152/jn.00651.2016.

Sacchet MD, LaPlante RA, Wan Q, Pritchett DL, Lee AKC, Hämäläinen M, Moore CI, Kerr CE, Jones SR. 2015. Attention Drives Synchronization of Alpha and Beta Rhythms between Right Inferior Frontal and Primary Sensory Neocortex. J Neurosci. 35(5):2074–2082. doi:10.1523/JNEUROSCI.1292-14.2015.

Scanziani M. 2000. GABA spillover activates postsynaptic GABAB receptors to control rhythmic hippocampal activity. Neuron. 25(3):673–681.

Scheyltjens, I., & Arckens, L. (2016). The current status of somatostatin-interneurons in inhibitory control of brain function and plasticity. Neural plasticity.

Sherman MA, Lee S, Law R, Haegens S, Thorn CA, Hämäläinen MS, Moore CI, Jones SR. 2016. Neural mechanisms of transient neocortical beta rhythms: Converging evidence from humans, computational modeling, monkeys, and mice. Proc Natl Acad Sci. 113(33):E4885–E4894. doi:10.1073/pnas.1604135113.

Sherman SM. 2016. Thalamus plays a central role in ongoing cortical functioning. Nat Neurosci. 19:533.

Shin H, Law R, Tsutsui S, Moore CI, Jones SR. 2017. The rate of transient beta frequency events predicts behavior across tasks and species. eLife. 6:e29086. doi:10.7554/eLife.29086.

Simon A, Oláh S, Molnár G, Szabadics J, Tamás G. 2005. Gap-Junctional Coupling between Neurogliaform Cells and Various Interneuron Types in the Neocortex. J Neurosci. 25(27):6278. doi:10.1523/JNEUROSCI.1431-05.2005.

Spitzer B, Haegens S. 2017. Beyond the Status Quo: A Role for Beta Oscillations in Endogenous Content (Re)Activation. eNeuro. 4(4):1–15. doi:10.1523/ENEURO.0170-17.2017.

Takahashi N, Oertner TG, Hegemann P, Larkum ME. 2016. Active cortical dendrites modulate perception. Science. 354(6319):1159–1165. doi:10.1126/science.aah6066.

Takahashi, N., Ebner, C., Sigl-Glöckner, J., Moberg, S., Nierwetberg, S., & Larkum, M. E. (2020). Active dendritic currents gate descending cortical outputs in perception. Nature Neuroscience, 23(10), 1277–1285.

Teleńczuk, B., Dehghani, N., Le Van Quyen, M., Cash, S. S., Halgren, E., Hatsopoulos, N. G., & Destexhe, A. (2017). Local field potentials primarily reflect inhibitory neuron activity in human and monkey cortex. Scientific reports, 7, 40211.

Tinkhauser G, Torrecillos F, Duclos Y, Tan H, Pogosyan A, Fischer P, Carron R, Welter M-L, Karachi C, Vandenberghe W, et al. 2018. Beta burst coupling across the motor circuit in Parkinson’s disease. Neurobiol Dis. 117:217–225. doi:10.1016/j.nbd.2018.06.007.

Kay, L. M., Beshel, J., Brea, J., Martin, C., Rojas-Líbano, D., & Kopell, N. (2009). Olfactory oscillations: the what, how and what for. Trends in neurosciences, 32(4), 207–214.

Torrecillos F, Alayrangues J, Kilavik BE, Malfait N. 2015. Distinct Modulations in Sensorimotor Postmovement and Foreperiod β-Band Activities Related to Error Salience Processing and Sensorimotor Adaptation. J Neurosci. 35(37):12753–12765. doi:10.1523/JNEUROSCI.1090-15.2015.

Vecchia, D., Beltramo, R., Vallone, F., Chéreau, R., Forli, A., Molano-Mazón, M., … & Fellin, T. (2020). Temporal Sharpening of Sensory Responses by Layer V in the Mouse Primary Somatosensory Cortex. Current Biology. 30(9):1589–1599.e10. doi: 10.1016/j.cub.2020.02.004

Viaene, A. N., Petrof, I., & Sherman, S. M. (2011). Synaptic properties of thalamic input to layers 2/3 and 4 of primary somatosensory and auditory cortices. Journal of neurophysiology, 105(1), 279–292.

Wessel, J. R. (2020). β-bursts reveal the trial-to-trial dynamics of movement initiation and cancellation. Journal of Neuroscience, 40(2), 411–423.

Westfall PH, Young SS. 1993. Resampling-based multiple testing: Examples and methods for p-value adjustment. John Wiley & Sons.

Williams LE, Holtmaat A. 2018. Higher-Order Thalamocortical Inputs Gate Synaptic Long-Term Potentiation via Disinhibition. Neuron. 101:91–102. doi:10.1016/j.neuron.2018.10.049.

Wimmer, V. C., Bruno, R. M., De Kock, C. P., Kuner, T., & Sakmann, B. (2010). Dimensions of a projection column and architecture of VPM and POm axons in rat vibrissal cortex. Cerebral cortex, 20(10), 2265–2276.

Wozny, C., & Williams, S. R. (2011). Specificity of synaptic connectivity between layer 1 inhibitory interneurons and layer 2/3 pyramidal neurons in the rat neocortex. Cerebral cortex, 21(8), 1818–1826.

Xing D, Yeh C-I, Shapley RM. 2009. Spatial Spread of the Local Field Potential and its Laminar Variation in Visual Cortex. J Neurosci. 29(37):11540–11549. doi:10.1523/JNEUROSCI.2573-09.2009.

Zanos S, Rembado I, Chen D, Fetz EE. 2018. Phase-Locked Stimulation during Cortical Beta Oscillations Produces Bidirectional Synaptic Plasticity in Awake Monkeys. Curr Biol. 28(16):2515–2526. doi:10.1016/j.cub.2018.07.009.

Zhang, W., & Bruno, R. M. (2019). High-order thalamic inputs to primary somatosensory cortex are stronger and longer lasting than cortical inputs. Elife, 8, e44158.

Ziegler DA, Pritchett DL, Hosseini-Varnamkhasti P, Corkin S, Hämäläinen M, Moore CI, Jones SR. 2010. Transformations in oscillatory activity and evoked responses in primary somatosensory cortex in middle age: A combined computational neural modeling and MEG study. Neuroimage. 52(3):897–912. doi:10.1016/j.neuroimage.2010.02.004.

