## Supplemental Materials for "Thalamocortical mechanisms regulating the relationship between transient beta events and human tactile perception"

**Supplementary Materials and Methods**

**Supplemental MEG Data Collection and Analysis Methods**

*Correlating Class 2 model waveforms and MEG miss trials*

To compare “Class 2” (in which a beta event precedes the tactile stimulus by ~100-400 ms) model waveforms to MEG miss trials, the mean empirical evoked response waveform (*u*_MEG_) was linearly regressed by least-squares directly onto the smoothed model waveform (x_model_), obtaining:

$$\mu_{MEG}\approx\beta_{0}+\beta_{1}\cdot x_{model}$$

The $r^{2}$ values assess agreement between model and data. We performed regressions separately for miss trials in event and no-event cases as well as in aggregate. In all cases, the constant offset parameter $\beta_{0}$ was small and not considered further. The linear scaling parameter $\beta_{1}$ was similar in all cases and used to estimate the scaling factor applied to the model to estimate the number of neurons that can account for the magnitude of the recorded signal. We report error regions reflecting standard error of the mean $\frac{s}{n^{1/2}}$.

*Classification of Class 1 evoked responses in MEG data*

We first calculated the normalized cross-correlation $\chi{(x}_{model},x_{MEG})$ between the model waveform and each MEG trial -- over the entire trial (-1000–2500ms) -- and found the lag time of peak cross-correlation for each trial i:

$$\tau_{i}=\text{argmax }\chi_{i}$$

We then restricted our analysis to candidate Class 1 trials whose peak cross-correlation occurred at lags between 0 and 25ms. This interval would not confound Class 1 with Class 3 trials, which exhibit similar waveforms but with different time-delays. This method coincidentally generated a sample small enough (n=18 trials) that all waveforms could be presented in one figure for inspection by the reader, but large enough that statistics could be meaningfully computed. Similar results were obtained in wider intervals, up to $-5ms\leq\tau\leq30ms$.

The model having indicated that beta phase at 25ms poststimulus should covary with detection, we then examined how this feature, in combination with cross-correlation, dissociated Class 1 candidates into hit and miss trials. To do so, we simply examined a circular histogram of beta phases for the 18 candidate Class 1 trials.

**Supplemental Computational Model Methods**

**Supplementary Table 1 (Model Parameters)**

The model consists of four cell types, of which only the two pyramidal cell types have multiple compartments and contribute to the dipole. Layer 2/3 pyramidal cells consist of a soma, an apical trunk directly above the soma, followed by apical 1 and, most distally, the apical tuft. The apical oblique dendrite branches horizontally from the apical trunk. There are also three basal dendritic compartments, with basal 1 directly below the soma and basal 2 and basal 3 branching from basal 1 at an angle, for a total of 8 compartments. The setup of the layer 5 pyramidal neurons is the same except that there is an additional apical compartment, apical 2, situated between apical 1 and the apical tuft. See Materials and Methods and Figure 1 for a diagram of cell morphologies. Figure 1 also provides a schematic of the connectivity between cell types in the model. Within each layer, synaptic strengths between pyramidal neurons are scaled according to a 2D symmetric Gaussian defined on the grid of cells, with a weight space constant of 3. Similarly, the synaptic delay between two PNs in the same layer is scaled according to an inverse Gaussian with a delay space constant of 3. Parameters describing the cellular properties, local network connectivity, evoked response, and beta event generation are provided in this table. Parameter changes from our prior modeling study Jones et al 2009 are highlighted and described further below in Supplemental Computational Model Methods.

| **Cell parameters** | | | |
| --- | --- | --- | --- |
| *Cell type* | *Parameter* | *Jones, et al 2009* | *Law, et al 2020* |
| L2/3 Pyr | Soma length (µm) | 22.1 | 22.1 |
|  | Soma diameter (µm) | 23.4 | 23.4 |
|  | Soma capacitive density (µF/cm^2^) | 0.6195 | 0.6195 |
|  | Soma resistivity (Ω-cm) | 200 | 200 |
|  | Dendrite capacitive density (µF/cm^2^) | 0.6195 | 0.6195 |
|  | Dendrite resistivity (Ω-cm) | 200 | 200 |
|  | Apical trunk length (µm) | 59.5 | 59.5 |
|  | Apical trunk diameter (µm) | 4.25 | 4.25 |
|  | Apical 1 length (µm) | 306 | 306 |
|  | Apical 1 diameter (µm) | 4.08 | 4.08 |
|  | Apical tuft length (µm) | 238 | 238 |
|  | Apical tuft diameter (µm) | 3.4 | 3.4 |
|  | Apical oblique length (µm) | 340 | 340 |
|  | Apical oblique diameter (µm) | 3.91 | 3.91 |
|  | Basal 1 length (µm) | 85 | 85 |
|  | Basal 1 diameter (µm) | 4.25 | 4.25 |
|  | Basal 2 length (µm) | 255 | 255 |
|  | Basal 2 diameter (µm) | 2.72 | 2.72 |
|  | Basal 3 length (µm) | 255 | 255 |
|  | Basal 3 diameter (µm) | 2.72 | 2.72 |
|  | AMPA reversal (mV) | 0 | 0 |
|  | AMPA rise time (ms) | 0.5 | 0.5 |
|  | AMPA decay time (ms) | 5 | 5 |
|  | NMDA reversal (mV) | 0 | 0 |
|  | NMDA rise time (ms) | 1 | 1 |
|  | NMDA decay time (ms) | 20 | 20 |
|  | GABA_A_ reversal (mV) | -80 | -80 |
|  | GABA_A_ rise time (ms) | 0.5 | 0.5 |
|  | GABA_A_ decay time (ms) | 5 | 5 |
|  | GABA_B_ reversal (mV) | -80 | -80 |
|  | GABA_B_ rise time (ms) | 1 | 45 |
|  | GABA_B_ decay time (ms) | 20 | 200 |
|  | Soma voltage-gated K^+^ channel density (S/cm^2^) | 0.01 | 0.01 |
|  | Soma Na^+^ channel density (S/cm^2^) | 0.18 | 0.18 |
|  | Soma leak channel reversal (mV) | -65 | -65 |
|  | Soma leak channel density (S/cm^2^) | 0.0000426 | 0.0000426 |
|  | Soma M-type K^+^ channel density (pS/µm^2^) | 250 | 250 |
|  | Dendrite voltage-gated K^+^ channel density (S/cm^2^) | 0.01 | 0.01 |
|  | Dendrite Na^+^ channel density (S/cm^2^) | 0.15 | 0.15 |
|  | Dendrite leak channel reversal (mV) | -65 | -65 |
|  | Dendrite leak channel density (S/cm^2^) | 0.0000426 | 0.0000426 |
|  | Dendrite M-type K^+^ channel density (pS/µm^2^) | 250 | 250 |
| L5 Pyr | Soma length (µm) | 39 | 39 |
|  | Soma diameter (µm) | 28.9 | 28.9 |
|  | Soma capacitive density (µF/cm^2^) | 0.85 | 0.85 |
|  | Soma resistivity (Ω-cm) | 200 | 200 |
|  | Dendrite capacitive density (µF/cm^2^) | 0.85 | 0.85 |
|  | Dendrite resistivity (Ω-cm) | 200 | 200 |
|  | Apical trunk length (µm) | 102 | 102 |
|  | Apical trunk diameter (µm) | 10.2 | 10.2 |
|  | Apical 1 length (µm) | 680 | 680 |
|  | Apical 1 diameter (µm) | 7.48 | 7.48 |
|  | Apical 2 length (µm) | 680 | 680 |
|  | Apical 2 diameter (µm) | 4.93 | 4.93 |
|  | Apical tuft length (µm) | 425 | 425 |
|  | Apical tuft diameter (µm) | 3.4 | 3.4 |
|  | Apical oblique length (µm) | 255 | 255 |
|  | Apical oblique diameter (µm) | 5.1 | 5.1 |
|  | Basal 1 length (µm) | 85 | 85 |
|  | Basal 1 diameter (µm) | 6.8 | 6.8 |
|  | Basal 2 length (µm) | 255 | 255 |
|  | Basal 2 diameter (µm) | 8.5 | 8.5 |
|  | Basal 3 length (µm) | 255 | 255 |
|  | Basal 3 diameter (µm) | 8.5 | 8.5 |
|  | AMPA reversal (mV) | 0 | 0 |
|  | AMPA rise time (ms) | 0.5 | 0.5 |
|  | AMPA decay time (ms) | 5 | 5 |
|  | NMDA reversal (mV) | 0 | 0 |
|  | NMDA rise time (ms) | 1 | 1 |
|  | NMDA decay time (ms) | 20 | 20 |
|  | GABA_A_ reversal (mV) | -80 | -80 |
|  | GABA_A_ rise time (ms) | 0.5 | 0.5 |
|  | GABA_A_ decay time (ms) | 5 | 5 |
|  | GABA_B_ reversal (mV) | -80 | -80 |
|  | GABA_B_ rise time (ms) | 1 | 45 |
|  | GABA_B_ decay time (ms) | 20 | 200 |
|  | Soma voltage-gated K^+^ channel density (S/cm^2^) | 0.01 | 0.01 |
|  | Soma Na^+^ channel density (S/cm^2^) | 0.16 | 0.16 |
|  | Soma leak channel reversal (mV) | -65 | -65 |
|  | Soma leak channel density (S/cm^2^) | 0.0000426 | 0.0000426 |
|  | Soma Ca^2+^ channel density (pS/µm^2^) | 60 | 0 |
|  | Soma Ca^2+^ decay time (ms) | 20 | 20 |
|  | Soma Ca^2+^-dependent K^+^ channel density (pS/µm^2^) | 0.0002 | 0.0002 |
|  | Soma M-type K^+^ channel density (pS/µm^2^) | 200 | 200 |
|  | Soma T-type Ca^2+^ channel density (S/cm^2^) | 0.0002 | 0.0002 |
|  | Soma HCN channel density (S/cm^2^) | 0.000001 | 0.000001 |
|  | Dendrite voltage-gated K^+^ channel density (S/cm^2^) | 0.01 | 0.01 |
|  | Dendrite Na^+^ channel density (S/cm^2^) | 0.14 | 0.14 |
|  | Dendrite leak channel reversal (mV) | -71 | -71 |
|  | Dendrite leak channel density (S/cm^2^) | 0.0000426 | 0.0000426 |
|  | Dendrite Ca^2+^ channel density (pS/µm^2^) | 60 | 60* |
|  | Dendrite Ca^2+^ decay time (ms) | 20 | 20 |
|  | Dendrite Ca^2+^-dependent K^+^ channel density (pS/µm^2^) | 0.0002 | 0.0002 |
|  | Dendrite M-type K^+^ channel density (pS/µm^2^) | 200 | 200 |
|  | Dendrite T-type Ca^2+^ channel density (S/cm^2^) | 0.0002 | 0.0002 |
|  | Dendrite HCN channel density (S/cm^2^) | 0.000001** | 0.00001** |

* Density is equal to this value in apical dendrites (apical trunk, apical oblique, apical 1, apical 2, and apical tuft) only, and equal to 0 in the basal dendrites (basal 1, basal 2, basal 3).

** HCN channel density increases as an exponential function of distance from the soma, starting at 0.000001 S/cm^2^ in the soma and increasing with a space constant of 0.003.

| **Local network connectivity parameters** | | | | |
| --- | --- | --- | --- | --- |
| *Source cell* | *Target cell* | *Synapse* | *Maximal conductance (µS)* | |
|  |  |  | *Jones 2009* | *Law 2019* |
| L2/3 Pyr | L2/3 Pyr | AMPA | 0.0005 | 0.0005 |
|  |  | NMDA | 0.0005 | 0.0005 |
|  | L2/3 Basket | AMPA | 0.0005 | 0.0005 |
|  | L5 Pyr | AMPA | 0.00025 | 0.00025 |
|  | L5 Basket | AMPA | 0.00025 | 0.00025 |
| L2/3 Basket | L2/3 Pyr | GABA_A_ | 0.05 | 0.05 |
|  |  | GABA_B_ | 0.05 | 0.05 |
|  | L2/3 Basket | GABA_A_ | 0.02 | 0.02 |
|  | L5 Pyr | GABA_A_ | 0.001 | 0 |
|  |  | GABA_B_ | 0 | 0.0002 |
| L5 Pyr | L5 Pyr | AMPA | 0.0005 | 0.0005 |
|  |  | NMDA | 0.0005 | 0.0004 |
|  | L5 Basket | AMPA | 0.0005 | 0.0005 |
| L5 Basket | L5 Pyr | GABA_A_ | 0.025 | 0.02 |
|  |  | GABA_B_ | 0.025 | 0.005 |
|  | L5 Basket | GABA_A_ | 0.02 | 0.02 |

| **Exogenous proximal and distal input parameters to generate the evoked response** | | | | |
| --- | --- | --- | --- | --- |
| *Input type* | *Target cell* | *Synapse* | *Maximal conductance (µS)* | |
|  |  |  | *Jones 2009* | *Law 2019* |
| Early proximal | L2/3 Pyr | AMPA | 0.001 | 0.0011 |
|  | L2/3 Basket | AMPA | 0.002 | 0.002 |
|  | L5 Pyr | AMPA | 0.0005 | 0.001 |
|  | L5 Basket | AMPA | 0.001 | 0.001 |
| Distal | L2/3 Pyr | AMPA | 0.001 | 0.004 |
|  |  | NMDA | 0.001 | 0.004 |
|  | L2/3 Basket | AMPA | 0.0005 | 0.0005 |
|  |  | NMDA | 0.0005 | 0.0005 |
|  | L5 Pyr | AMPA | 0.001 | 0.0005 |
|  |  | NMDA | 0.001 | 0.0005 |
| Late proximal | L2/3 Pyr | AMPA | 0.0053 | 0.005 |
|  | L2/3 Basket | AMPA | 0.0053 | 0.005 |
|  | L5 Pyr | AMPA | 0.0027 | 0.01 |
|  | L5 Basket | AMPA | 0.0027 | 0.01 |

| **Beta event parameters** | | | | |
| --- | --- | --- | --- | --- |
| *Burst properties* | | | | |
| *Input type* | *Parameter* | | *Sherman 2016* | *Law 2019* |
| Proximal | Standard deviation (ms) | | 20 | 20 |
|  | Number of bursts | | 10 | 10 |
|  | Spikes per burst | | 2 | 2 |
|  | L2/3 delay (ms) | | 0.1 | 0.1 |
|  | L5 delay (ms) | | 1.0 | 1.0 |
| Distal | Standard deviation (ms) | | 10 | 10 |
|  | Number of bursts | | 10 | 10 |
|  | Spikes per burst | | 2 | 2 |
|  | L2/3 delay (ms) | | 5 | 0.5 |
|  | L5 delay (ms) | | 5 | 0.5 |
| *AMPA conductances (µS)* | | | | |
| *Input type* | *Target cell* | *Compartment* | *Sherman 2016* | *Law 2019* |
| Proximal | L2/3 Pyr | Basal 2 | 0.00002 | 0.00002 |
|  |  | Basal 3 | 0.00002 | 0.00002 |
|  |  | Apical oblique | 0.00002 | 0.00002 |
|  | L2/3 Basket | Soma | 0.00004 | 0.00004 |
|  | L5 Pyr | Basal 2 | 0.00002 | 0.00002 |
|  |  | Basal 3 | 0.00002 | 0.00002 |
|  |  | Apical oblique | 0.00002 | 0.00002 |
|  | L5 Basket | Soma | 0.00002 | 0.00002 |
| Distal | L2/3 Pyr | Apical tuft | 0.00004 | 0.00008 |
|  | L2/3 Basket | Soma | 0.00008 | 0.00032 |
|  | L5 Pyr | Apical tuft | 0.00004 | 0.00004 |

*Comparison between current model and Jones et al. 2009*

Supplemental table 1 highlights differences in parameters between the Jones et al 2009 model and the model used here. There were also some network structure differences between the 2009 model and current model. In the current model, we added a Martinotti-like recurrent tuft connections from the L5 interneuron to the L5 pyramidal neurons’ distal dendrites and removed the L2/3 GABA_A_ interneuron synapses onto those same dendrites. We also adapted the model’s L5 calcium channel distribution, restricting expression to the apical dendrite. The most important *a priori* change in the model was an increase of the GABA_B_ channel time constants to reflect data reported in (Otis et al. 1993). We reduced this model to a double-exponential conductance with 45/200ms respective rise/fall times, matching the peak conductance latency ~100ms of the above channel model (see also Figure 1 for model details). Of note, the default network model currently distributed with the Human Neocortical Neurosolver software (Neymotin et. al, 2020) reflects the network structure in Jones et al 2009.

*Modeling the tactile evoked response*

Model simulations of tactile evoked responses were generated by a sequence of three inputs to SI, as in (Jones et al. 2007; Jones et al. 2009). Based on this prior work, the first “feedforward”, or bottom-up, input is simulated to arrive at pyramidal basal dendrites 25ms after stimulus onset, ostensibly from L4 by way of the sensory thalamus. Then, at 70ms, a top-down “feedback” input arrives from SII or higher-order thalamus. A subsequent basal input from L4 arrives 135ms poststimulus; however, we restricted our analysis of evoked responses to the first 140ms following the tactile stimulus because variability in the late input may depend on unmodeled interactions between SI and other cortical regions e.g. premotor cortex (Auksztulewicz and Blankenburg 2013). See Supplementary Table 1 for parameters of evoked inputs.

We obtained substantially better agreement with evoked response curvature by smoothing the raw model evoked response with a 45ms Hamming window. A primary mismatch between model and data was the M70 depth, which is shallow in the model compared to the data. Our primary conclusions hold true irrespective of this discrepancy.

*Dependence of model beta event dipole on slow inhibitory conductances*

A key difference between the present model and Sherman et al., 2016 is that the additional slow inhibition can affect the event dipole itself, yielding at least some beta events that couple to lower-frequency (delta/theta) bands (Supplementary Figures 1D/E and Supplementary Figure 2, cf. Carracedo et al, 2013). More specifically, the beta event waveform shape was largely maintained, however GABA_B_ slowly hyperpolarizes L2/3 somatic and L5 apical compartments with opposing effects, leading to downward dipoles from L2/3 and upward dipoles from L5. The latter has a dominant effect on the net current dipole, leading to an upward deflecting signal and a longer dipole tail after the beta event (see 250-500ms in Figure 5C, and Supplementary Figure 1D/E). In the spectral domain, this tail generates increased power in the theta/delta bands, but the power is none-the-less primarily concentrated near 20Hz as in our data .

The model beta event’s waveshape depends, at least in part, on the relative strength of slow inhibitory coupling in L2/3 somata and L5 apical dendrites. Supplementary Figure 1 demonstrates this fact by simulating a beta events for several GABA_B_ conductances in the L5 mid-apical dendrite, with the L2/3 somatic conductance held fixed. For larger L5 apical GABA_B_ conductances, the events are highly asymmetric and as the inhibition is increased high power theta frequency activity emerges (see Supplementary Figure 1D/E) and coupling in the theta/delta bands is apparent.

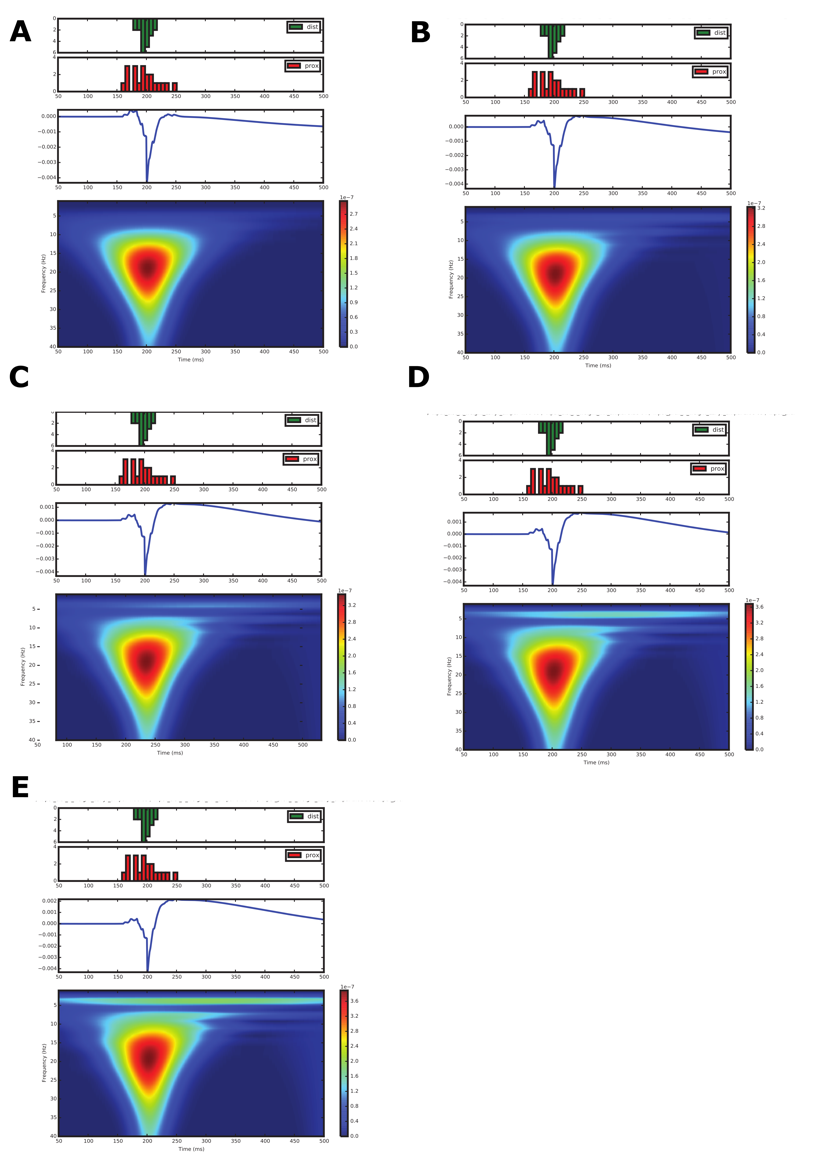

**Supplementary Figure 1: Beta event waveshapes vary with slow inhibitory conductances in the L5 apical dendrite.** Model events are shown (as per Figure 5B). Model GABA_B_ conductances in the L5 dendrite are **A)** 1pS **B)** 2pS **C)** 3pS **D)** 4pS and **E)** 5pS. The model beta event in the main text corresponds to (B).

*Existence of asymmetric beta events in MEG*

Given the results in Supplementary Figure 1, we might expect a variety of beta-event waveshapes in our MEG data depending on the local cortical expression of GABA_B_ receptors. This may appear to be at odds with the characterized left/right event symmetry reported in (Sherman, et al 2016). That study, however, analyzed only the 50 most powerful beta events. In our study, we have considered a much larger number of events (see main text). Indeed, Supplementary Figure 2 shows an example of an empirically determined asymmetric beta event, found by direct cross-correlation of a model event with MEG data.

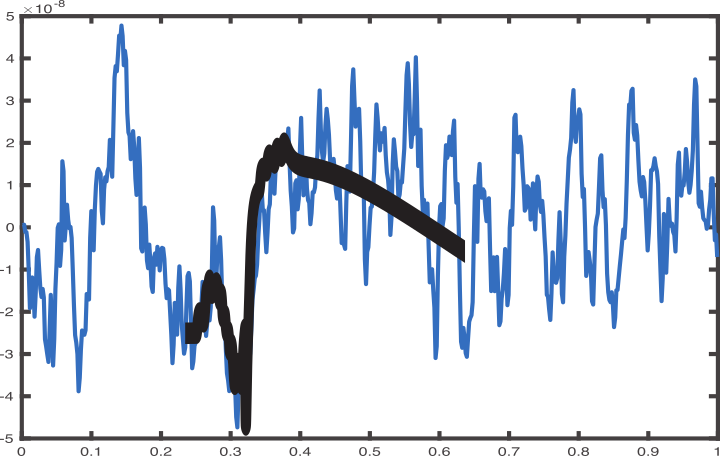

**Supplementary Figure 2: Existence of asymmetric beta events in MEG data.** Example of MEG beta event (blue) consistent (i.e., high cross-correlation value) with asymmetric the model beta event during GABA_B_ inhibition in the

*Robustness of the SI evoked response results to noise*

In the main text, we modeled evoked responses in the absence of background activity, aside from those inputs corresponding to the beta event itself, assuming the influence of high-power macroscale beta events have a dominant impact on the evoked response. Here, we introduce a small amount of subthreshold background noise simulated as noisy excitatory (AMPA) synaptic drive, whose time of activation follows a Poisson process, arriving at the basal dendrite due to e.g. spontaneous layer 4 spiking activity. The amplitude of this noise is consistent with the amplitude of noise observed around the high-power beta events in our data (green box Figure 2A) and present during the entire simulation. The model generates three latency-dependent classes of evoked responses, reproducing the same effect in the model, as without background noise (Supplementary Figure 3A,B,C,D, compare to Figure 6). In general, the main results are robust to noise, unless it is strong enough to make the pyramidal neurons spike before stimulus onset.

Importantly, all simulations presented are generated from a network of fixed size (100 PN per layer). The amplitude of the simulated subthreshold (i.e., no spiking in the pyramidal neurons) beta events in this fixed sized network is an order of magnitude smaller than the simulated evoked response, in which significant pyramidal neuron spiking occurs (compare dipole amplitudes in Supplementary Figure 3A and 3B). This comparison is emphasized by blowing up the early Class 3 dipole response from single pyramidal neuron spike (magenta in the spike histogram), which is an order of magnitude larger than the surrounding subthreshold activity in Figure 3E. Here, and in prior reports (Jones et al 2009, Sherman et al 2016), we presume that high-power beta events emerge over a significantly larger *subthreshold* network than the *spiking* network that contributes to the evoked response (Figure 3F). This is consistent with fact that in the recorded MEG data beta events amplitudes are an order magnitude greater (Figure 2A top, ~100nAm) than tactile evoked response amplitudes (Figure 3, ~10nAm). This is accounted for in our model by applying a larger scaling factor when comparing simulating beta events to data (not shown here, see Sherman et al 2016), than when comparing simulating evoked responses to data (e.g. Figure 9).

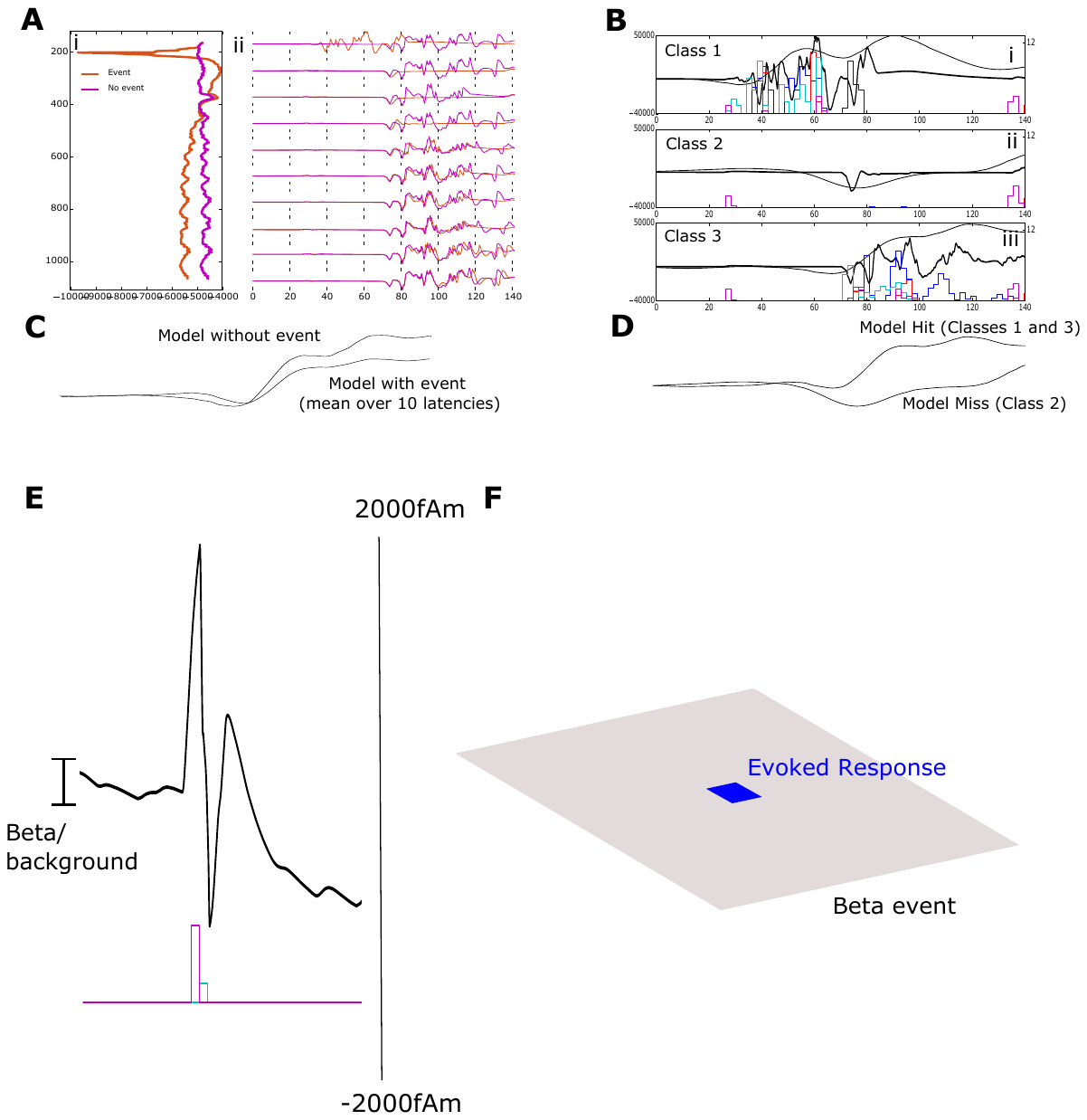

**Supplementary Figure 3:** **Model evoked responses in the presence of background activity**. **A)** Model evoked responses vs. beta event latency (compare to main text Figure 6) i) ongoing beta event with or without additional background activity (x-axis current dipole, y-axis time). ii) Model evoked responses (x-axis time, y-axis current dipole, *vice versa* from (i). **B)** Model class-average waveforms (compare to main text Figure 7) **C)** Mean event vs. no-event evoked responses, averaged over latencies. **D)** Mean Class 1 and 3 detected and Class 2 nondetected responses. **E)** Zoomed-in early Class 3 response (0—50ms), showing that the dipole amplitude from even one pyramidal spike (magenta) and its attendant inhibitory activity (cyan) is significantly larger the surrounding subthreshold beta and/or noise driven activity. F) Schematization of the relative cortical surface sizes contributing to beta events and early evoked responses: beta events in the MEG data are higher-power than evoked responses because they are generated over a significantly larger network. However, in our fixed size model simulations subthreshold beta events and noise are lower in amplitude than evoked responses (A and B), a difference that is accounted for with different model scaling factor when comparing directly to MEG data.

*Time-Derivative corresponding to Figure 6*

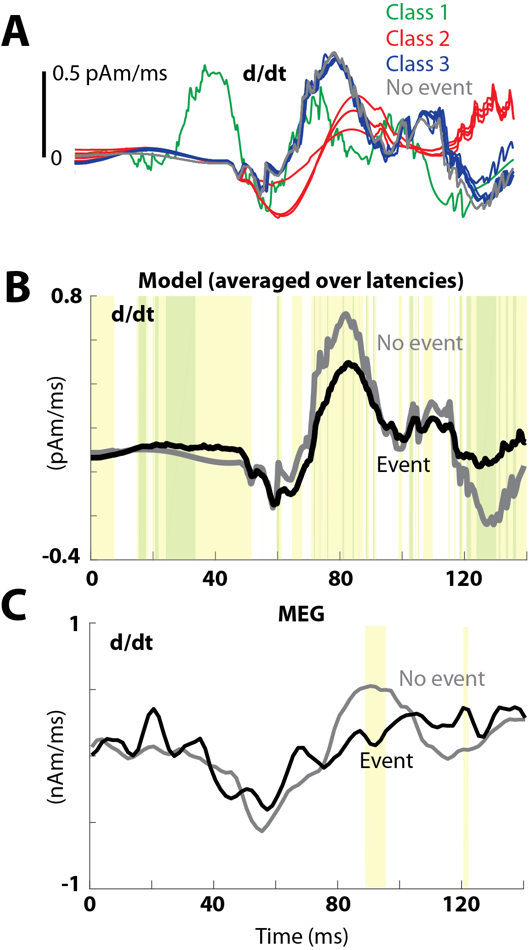

**Supplementary Figure 4: Time-derivatives corresponding to Figure 6.** **A.** Time derivative of each of the evoked response simulations as shown in Figure 6E color coded for Class 1 (green), Class 2 (red), Class 3 (blue) and no event trials (gray). **B.** Time-derivatives of the simulated evoked responses averaged over all Classes (black) compared to averaged no event evoked response (gray), corresponding to Figure 6A. **C**. Time-derivates of MEG recorded evoked responses averaged of event (black) and no event trials (gray), corresponding to Figure 6B.
